# A modular model of immune response as a computational platform to investigate a pathogenesis of infection disease

**DOI:** 10.1101/2024.08.19.608570

**Authors:** M.I. Miroshnichenko, F.A. Kolpakov, I.R. Akberdin

**Affiliations:** Department of Computational Biology, Scientific Center for Genetics and Life Sciences, Sirius University of Science and Technology, Sirius, Russia

**Keywords:** coronavirus, SARS-CoV-2, COVID-19, mathematical model, BioUML, immune response

## Abstract

The COVID-19 pandemic significantly transformed the field of mathematical modeling in immunology. International collaboration among numerous research groups yielded a substantial amount of experimental data, which greatly facilitated model validation and led to the development of new mathematical models. The aim of the study is an improvement of system understanding of the immune response to SARS-CoV-2 infection based on the development of a modular mathematical model which provides a foundation for further research on host-pathogen interactions. We utilized the open-source BioUML platform to develop a model using ordinary, delay and stochastic differential equations. The model was validated using experimental data from middle-aged individuals with moderate COVID-19 progression, including measurements of viral load, antibodies, CD4+ and CD8+ T cells, and interleukin-6 levels. Parameter optimization and sensitivity analysis were conducted to refine the model’s accuracy. The model effectively reproduces moderate, severe, and critical COVID-19 progressions, consistent with experimental observations. We investigated the efficiency and contributions of innate and adaptive immunity in response to SARS-CoV-2 infection and assessed immune system behavior during co-infection with HIV and organ transplantation. Additionally, we studied therapy methods, such as interferon administration. The developed model can be employed as a framework for simulating other infectious diseases taking into account follow-up immune response.

**Author summary:** Despite the significant progress reached in understanding of COVID-19, traditional methods still struggle to analyze and interpret the extensive and sometimes controversial experimental data on SARS-CoV-2 infection. Mathematical and systems biology approaches attempt to address this challenge by developing mathematical models of the immune response. We aimed not only to investigate the disease at a systemic level but also to provide a framework for further research on host-pathogen interactions, both existing and forthcoming. To achieve this, we constructed a model incorporating both innate and adaptive immunity, as well as cellular and humoral components. This together allowed us to conduct a series of *in silico* experiments, exploring the immune response across various levels and compartments. The results of these investigations offer valuable insights into the complex dynamics of the immune system and can guide future research and therapeutic strategies.

## 1. Introduction

The emergence of severe acute respiratory syndrome coronavirus 2 (SARS-CoV-2) led to one of the largest pandemics in human history, comparable with the Spanish flu and HIV/AIDS in terms of case numbers. SARS-CoV-2 causes Coronavirus disease 2019 (COVID-19) and is classified as a positive-sense single-stranded RNA virus [1] that infects human cells via the angiotensin-converting enzyme 2 (ACE2). It is a common membrane receptor expressed in most tissues and organs, primarily including the gastrointestinal tract, upper and lower airways, and circulatory system. Additionally, it is present in the brain, kidneys, testis, and other organs [2]. Such a wide diversity of target organs leads to an intricate and diverse disease progression with significant variability in severity. According to the National Institutes of Health, COVID-19 manifestations are categorized into five groups: asymptomatic infection, mild, moderate, severe, and critical conditions [3]. The distinction between mild and severe forms includes the progression of the virus to the lower respiratory tract and a decline in oxygen saturation level. In more severe cases, this leads to more serious symptoms, which, in the most critical cases, may result in respiratory failure and multiple organ dysfunction. The severity of COVID-19 depends on many factors, with age and chronic conditions such as asthma, cancer or diabetes being among the most significant [4]. It has been shown that the efficiency and quantity of various arms of the immune system diminish with age. These changes include a decrease in the activity of both neutralizing and non-neutralizing antibodies [5], as well as a reduction in the number of circulating T cells, particularly memory and differentiated effector T cells. Additionally, there is a decline in the proportion of T cells capable of producing cytotoxic molecules such as granzymes and perforins, which lead to impaired effector functions of T cells [6]. Similar processes also affect B cell function [7,8]. All of these factors are discussed in detail in Section 3.3.

Mathematical modeling is a widely used approach for studying various infections and following immune responses. It not only facilitates the exploration of the intricate behavior of distinct immune components during infection but also provides the opportunity to consider the unique biological characteristics inherent to different pathogens. While numerous mathematical models have been developed for various infectious agents, those affecting human health the most, such as influenza virus or HIV, are particularly prevalent. This is evident from the numerous models created in recent decades [9–14]. The surge in the number of models has been facilitated by the growing availability and accessibility of experimental data, along with the opportunity to use previously developed models as the basis for new ones.

There is a wide variability in the complexity of existing mathematical models, ranging from simplified ones with only a few ordinary differential equations [14,15] to more complex models composed of dozens or even multiple reactions [11]. This diversity provides an opportunity to examine the immune response and infection process from multiple perspectives. As an example, Hancioglu with co-authors [10] investigated the effects of drugs, including interferons, and initial viral load on influenza disease progression. In contrast, Heldt and co-authors [11] delved into the intracellular aspects of influenza, providing a detailed description of replication, transcription, and translation processes.

With the onset of the COVID-19 pandemic, there has arisen a demand to develop mechanistic models that describe the dynamics of SARS-CoV-2 in the human body, taking into account the subsequent immune response to the infection. Most studies published so far concentrate on SARS-CoV-2 infecting only the epithelial cells of the human lungs. For instance, Li and co-authors [16] built a low-dimensional model of coronavirus replication in alveolocytes, comprising only healthy and infected cells along with viral particles. Another simplified model by Du and Yuan [17] also examines SARS-CoV-2 but from the perspective of T-cell and antibody responses. In contrast, the more complex model developed by Leander and co-authors [12] encompasses not only interactions between the virus and epithelial cells in the lungs but also the reactions of the innate immunity, involving macrophages, chemokines and interferons. A notable feature of this model is its detailed classification of epithelial cells into five groups based on cell types, ACE2 receptor characteristics, and susceptibility to interferons.

Meanwhile, other researchers have explored SARS-CoV-2 infection not just from the perspective of cell interactions but also by considering biochemical processes, as demonstrated in the model by Wang and co-authors [18]. They focused on the relationships between intracellular, intercellular and organism levels, allowing for insights into how the behavior of the entire system changes under varying conditions at each level. Beyond that, the authors modeled various disease progressions, encompassing asymptomatic, mild-moderate, and severe modes. Furthermore, this model also considers significant features such as heterogeneity in the number of receptors on the cell surface and exhaustion of T cells. Additionally, it incorporates the implementation of distinct treatment strategies, including accelerating the interferon response or inhibiting T cell exhaustion. The study found that a robust interferon response is crucial for controlling the onset of symptoms. Besides it, the authors confirmed that T cell depletion significantly contributes to disease progression towards severe states, often accompanied by a cytokine storm. Finally, the study demonstrated that combination therapy proved itself to be the most effective treatment strategy for severe patients.

There are also unique models, such as Grebennikov’s SARS-CoV-2 model [19], which provides a detailed analysis of the virus’ intracellular life cycle kinetics. This model covers all major steps in coronavirus entry and replication, even considering specific processes such as uncoating and virion assembly. With a simulation spanning 24 hours, which approximately corresponds to a single replication cycle of SARS-CoV-2, the model allows for a detailed examination of each reaction. Being a high-resolution model, it has numerous applications for predicting potential targets for antiviral therapy. The authors identified key parameters, such as RNA degradation and the translation of non-structural proteins that significantly influence the virus life cycle. It is important to note that this model can be integrated into a multiscale mathematical model of SARS-CoV-2 infection, providing valuable insights into the intracellular dynamics of the virus.

Another well-formulated model of the SARS-CoV-2 infection process was developed by Grebennikov and colleagues [20]. The model integrates both innate and adaptive immune responses, incorporating plasma cells and antibody formation, along with the development of cytotoxic T cells. It specifically categorizes epithelial cells into infected, interferon-protected, and damaged types that facilitate a detailed examination of SARS-CoV-2 infection progression. Furthermore, the authors distinguished between antigen-presenting dendritic cells and interferon-producing cells. As a result, they identified thresholds for the increase of parameters related to innate and adaptive responses that contribute to the prolonged persistence of SARS-CoV-2. One of the key findings highlighted is that the coordinated kinetics of innate and adaptive responses is crucial for the optimal induction of the entire immune cascade.

One of the most complex models of the COVID-19 infection and immune response developed to date is by Zhou and colleagues [21]. It is composed of thirty-two ordinary differential equations and incorporates key immune system components such as neutrophils, natural killer cells, and most importantly, T regulatory cells which are often overlooked in other models but are crucial for orchestrating the immune response. It also integrates mucosal immunity and accounts for variations in SARS-CoV-2, including spike protein mutations, viral affinity for the ACE2 receptor, and replication efficiency. The authors confirmed the significance of IL-6 as a biomarker for severe symptoms and highlighted differences in immune response between patients with mild and severe symptoms.

Mathematical models have proven to be effective tools for studying diseases, addressing not only epidemiological aspects but also cell development and their interactions during immune response, intracellular pathogen dynamics, and specific scenarios like airborne transmission. The significance of modeling was especially evident during the COVID-19 pandemic. The surge in research produced a vast amount of diverse and often controversial experimental data, which traditional methods struggled to analyze and interpret. Mathematical modeling and a systematic approach were essential in handling this complexity. We estimated that over 40 mathematical models related to COVID-19 exist, primarily sourced from the BioModels web repository [22] and through manual searches, demonstrating the high demand for such models. Even after the end of the pandemic, the World Health Organization continues to report registered COVID-19 cases in its weekly epidemiological updates for the last few years [23], thereby demonstrating the residual perturbation of the virus in the population. This persistent presence suggests potential seasonality and possible outbreaks, similar to influenza, and raises concerns about long COVID [24]. Overall, these factors underscore the necessity for ongoing research in systems biology to deepen our understanding of SARS-CoV-2 infection and enhance our readiness for future pandemics.

Our research focuses on the detailed mechanisms of immune response development, encompassing both innate and adaptive arms, and considers cellular and humoral components. We pay close attention to the experimental data used for model calibration to ensure the reliability of results and predictions. Additionally, we addressed the impact of immune system aging on immune response efficiency. The modular model we developed is employed to identify potential targets for antiviral therapy and to verify a set of biological hypotheses. The model can also be utilized for fundamental studies on virus replication and the resulting immune response. Moreover, it may also serve as a framework for modeling infections caused by other viral pathogens, as its structure reflects common biological processes underlying the classical antiviral immune response.

## 2. Materials and methods

### 2.1 Mathematical model

To construct a complex multi-scale mathematical model of the immune response to SARS-CoV-2 infection, we employed a modular approach previously that had been used for building various mathematical models [25,26], along with model manual extension and adjustment. At the first step we integrated manually reviewed and validated models of the immune response to *Mycobacterium tuberculosis* (MT) [27] and Influenza A (IA) virus [13] infections. Both models are two-compartmental and include the lungs and draining lymph nodes as separate compartments. The MT model focuses on the innate immune response, highlighting the roles of antigen-presenting cells and T helper cells, while the IA model predominantly emphasizes adaptive immunity, providing a detailed description of both humoral and cellular components.

The MT model, developed by Marino and Kirschner, describes the infection of resting macrophages (*M*_*R*_) in the lungs by bacteria (*B*_*T*_), which are categorized into external (*B*_*E*_) and internal (*B*_*I*_) types. The bacteria trigger the activation of immature dendritic cells (*IDC*) and resting macrophages, leading to their differentiation into mature dendritic cells (*MDC*) and activated macrophages (*M*_*A*_), respectively. MDCs then drive the activation of naïve T cells (*T*_*naive*_) to form precursors (*T*_*P*_) in the lymph nodes. It should be noted that T cell species combine both cytotoxic and T helper functions. In the lungs, T cell precursors differentiate into T helper 1 and 2 types (*Th*_1_, *Th*_2_), which orchestrate cytokine production and contribute to the elimination of infected macrophages. At the same time, cytokines act as regulatory factors for the vast majority of immune response reactions.

The IA model, built by Lee and co-authors, simulates the infection of epithelial cells (*E*_*P*_) in the lungs by the influenza virus (*V*). Infected epithelial cells (*EP*_*I*_) begin to produce viral particles, which activate *IDC* leading to their transport to the lymph nodes, where they mature into mature dendritic cells (*MDC*). They drive the activation of naïve T helper cells (*H*_*N*_), naïve cytotoxic T cells (*T*_*N*_), and naïve B cells (*B*_*N*_), resulting in their proliferation into effector cells (*H*_*E*_, *T*_*E*_ and *B*_*A*_ respectively). Effector T helper cells regulate the differentiation of B cells into short-lived (*P*_*S*_) and long-lived (*P*_*L*_) plasma cells. The latter produce antibodies (*A*) that eliminate free viral particles in the lungs. Eventually, effector cytotoxic T cells migrate to the lungs and eradicate infected epithelial cells.

Based on the mentioned models, we have developed an extended model comprising three compartments — the upper airways, lungs and lymph nodes, representing the key areas involved in respiratory infection (Figure 1). Complete and detailed model description is given in S1 Text. The constructed modular model is also available on GitLab: https://gitlab.sirius-web.org/diploms/modular-immune-system. To address the common issue of model reproducibility in systems biology [28], we additionally reproduced the model using Julia programming language and conducted all equivalent investigations. The Julia version is also available on GitLab.

**Figure 1.**
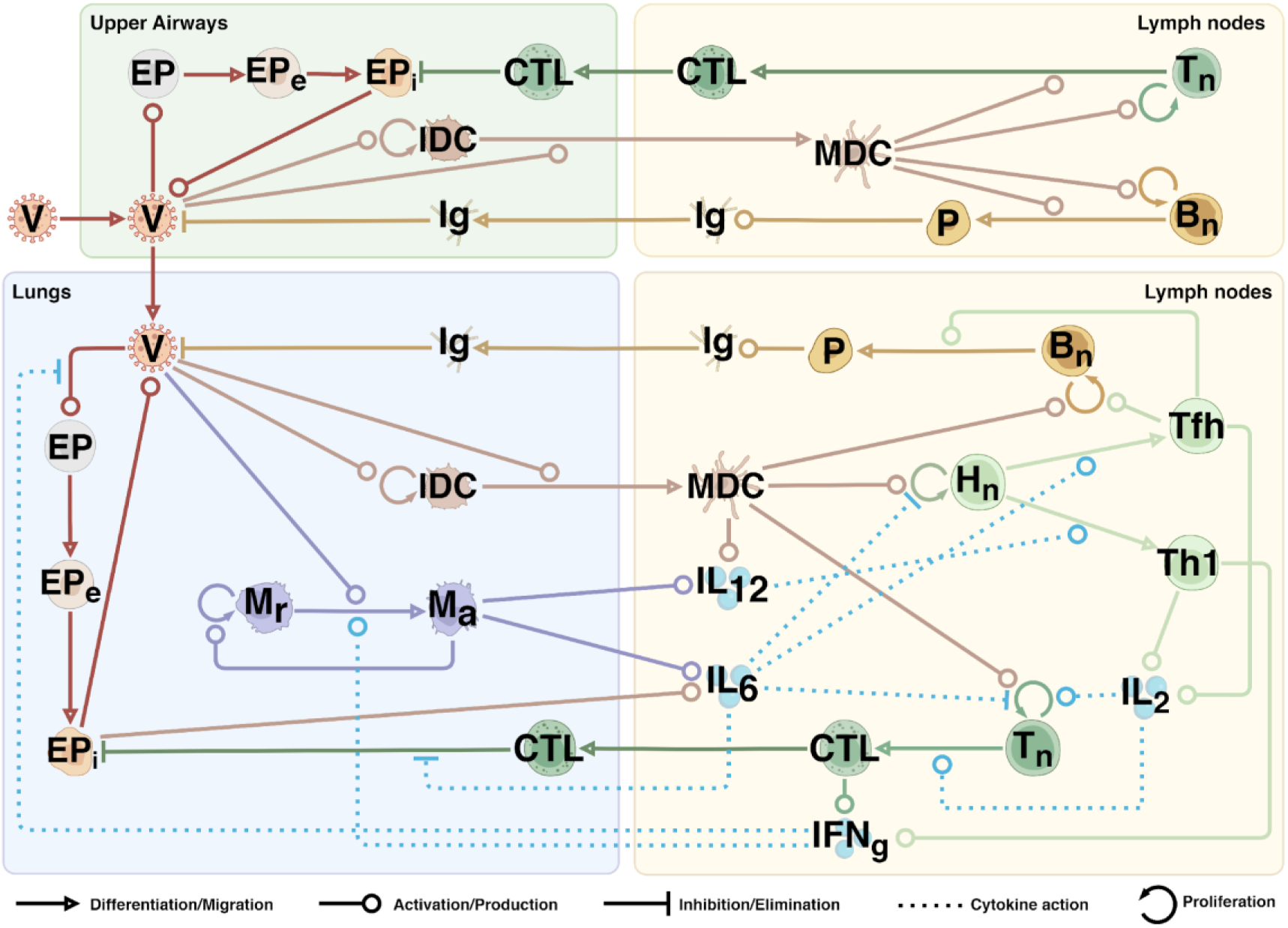
A schematic representation of the built model. The process notations are provided below the diagram. V – virus, EP – epithelial cells (e – exposed, i – infected), DC – dendritic cells (I – immature, M - mature), M – macrophages (r – resting, a – activated), Hn – naïve CD4+ T cells, Tn – naïve CD8+ T cells, Bn – naïve B cells, Th1 – T helper 1 type cells, Tfh – T follicular helper cells, CTL – cytotoxic T cells, P – plasma cells, Ig – immunoglobulins (IgA, IgM, IgG), IL – interleukins (2, 6, 12), IFNg – interferon gamma.

### 2.2 Experimental data

To accurately reflect biological reality and successfully verify the model, we used various experimental data, including time-series dynamics of cells and molecules, calculated or estimated parameters, as well as the initial values for model entities. Some parameter values were taken from existing mathematical models [12,13,19–21,27], while others were calculated independently.

#### 2.2.1 Initial values

To calculate the number of epithelial cells, we followed two assumptions. First, we considered only those cells which are susceptible to the infection and simultaneously express both ACE2 and TMPRSS2 receptors [29,30] required for coronavirus entry. Although the exact distribution of these receptors is difficult to determine, approximate data suggest that only about 1% of cells co-express both receptors and may therefore be infected [31]. Second, we based our estimates on the spatial distribution of cells. We initially assessed the epithelial surface area of the upper airways, including the nasal and oral cavities and the pharynx. The approximate surface areas are as follows: nasal mucosa – 170 cm^2^ (ranging from 150 to 200 cm^2^) [32], oral mucosa – 200 cm^2^ (ranging from 175 to 225 cm^2^) [33,34], and pharynx, which is composed of the oropharynx, nasopharynx, and hypopharynx – 150 cm^2^.

Since complete measurements for the pharynx are unavailable, we roughly estimated its surface area by considering the pharynx as a cylinder without bases. With an approximate length of 12 – 14 cm [35] and a lumen diameter of 2 cm [36,37], the pharynx has an estimated surface area of 150 cm^2^. Calculations for individual parts of the pharynx, such as the nasopharynx, verify this estimate and align with experimental data. In particular, the nasopharynx has a length of 4 cm and a lumen diameter of 2 cm [37], giving an area of about 50 cm^2^, which corresponds to known measurements [38].

Overall, the upper airways have an approximate surface area of 50000 mm^2^. Given that the density typical for airway epithelium ciliated cells is roughly 7,000 cells per mm^2^ [39,40], and considering only susceptible cells, we estimate the number of epithelial cells to be around 5.2 × 10^6^ cells. This result is consistent with a similar calculation using the experimentally derived density of susceptible cells in the airway epithelium, approximately 100 cells per mm^2^ [41], which yields about 5.5 × 10^6^ cells. Due to the difficulties in precise measurements, we opted for the higher estimate and used the initial count of 5.5 × 10^6^ susceptible epithelial cells in the upper airways.

We considered the trachea, bronchial tree, and lungs as the lower airways. According to cell counts provided by Hatton and colleagues [42], the approximate total number of ciliated, goblet, and alveolar (both AT1 and AT2) epithelium cells in the lower airways ranges from 3 × 10^10^ to 7 × 10^10^. Taking into account the proportion of vulnerable cells [31], we estimated the initial number of susceptible cells in the lower airways to be 5.5 × 10^8^. We considered both type I (AT1) and type II (AT2) alveolocytes due to emerging evidence about SARS-CoV-2 cell tropism to both of them, which was not clearly understood early in the pandemic [43,44]. Furthermore, we did not account for variability in ACE2 receptor expression, which is influenced by cell type, location, and developmental stage. Although this may potentially affect COVID-19 progression and severity to some extent [45,46].

To determine the initial number of dendritic cells (DCs), we used the known density of DCs in the airway epithelium, which varies widely from roughly 500 to 800 cells per mm^2^ of the epithelium [47–49]. We opted for the lower bound because several subpopulations of DCs are usually distinguished [50], yet the main antigen-presenting function is performed primarily by myeloid dendritic cells [51]. Given the surface area of the airway epithelium is around 50000 mm^2^, we calculated the total number of dendritic cells residing in the epithelium to be about 2.5 × 10^7^. To find the concentration of DCs, we first estimated the total volume of the epithelium in the upper airways. With an epithelial thickness ranging from 50 to 300 μm, and a mean value of 200 μm [52–54], the total volume is approximately 10 ml. This results in a dendritic cell concentration in the upper airways of 2.5 × 10^6^ cells per ml.

Due to the limited understanding of dendritic cell concentrations in the lungs, we performed calculations similar to those used for the upper airways. The lung surface area (excluding the bronchi) is approximately 70 m^2^, or 7 × 10^7^ mm^2^ [55]. The surface areas of the bronchi and trachea are 2.5 × 10^5^ mm^2^ and 2.4 × 10^5^ mm^2^ respectively. Assuming an average alveolar thickness of 0.3 μm [56] and a mean thickness of 50 μm [53,54,57] for the bronchi and trachea, we estimated the total lung volume to be 50000 ml. Given that myeloid DCs (mDCs) are the predominant subpopulation in the lungs [50], we opted for the upper bound density of 800 cells per mm^2^ of the epithelium [47–49]. Thus, we estimated the total number of DCs to be approximately 6 × 10^10^ cells, resulting in their concentration in the lungs of 1.2 × 10^6^ cells per ml.

Given that the known total number of macrophages in the lungs is 2 × 10^10^ cells [42] and taking into account the volume of the lower airway epithelium is 50000 ml, we estimated the concentration of alveolar macrophages to be 4 × 10^5^ cells per ml. It is worth noting that the estimated total number of macrophages is also supported by calculations based on a lung mucosal surface area of 100 m^2^, resulting in approximately 1.2 × 10^10^ macrophages, assuming their mean density of 200 cells per mm^2^ [58,59].

To accurately determine the initial concentrations of T and B cells, it is essential to consider the lymph nodes, which play a crucial role as reservoirs for lymphocytes. First, we evaluated the volume of the lymph nodes draining the upper and lower airways. For the upper airways, which include the head and cervical lymph nodes, their number ranges from 300 to 400 nodes [60,61], with an average volume of 0.3 ml per node [62,63]. This results in a total volume of lymph nodes in the upper airways ranging from 90 to 120 ml. We opted for 90 ml, based on other research indicating a lower mean volume for these nodes [64]. In contrast, the number of lymph nodes draining the lower airways [65], including the lungs, is much smaller and equals approximately 100 nodes [66]. However, they are larger, with an average volume of about 2 ml each [67]. Thus, we estimated the total volume of the lymph nodes draining the lower airways to be around 200 ml.

To evaluate the initial concentrations of naive B and T cells, we used the total number of lymphocytes in the lymph nodes [42] along with the estimated volumes of these nodes. For the lymph nodes in the upper airways, with a total volume of 90 ml, we calculated the concentrations of naive B cells and CD8+ T cells to be 3.3 × 10^4^ cells/ml and 1.6 × 10^4^ cells/ml correspondingly. In contrast, for the lymph nodes draining the lower airways, which have an estimated volume of 200 ml, the concentrations were found to be 6 × 10^4^ cells/ml for naïve B cells, 1.0 × 10^5^ cells/ml for naïve CD4+ T cells, and 3.3 × 10^4^ cells/ml for naïve CD8+ T cells. It should be noted that these calculations are based on the proportion of specific naive lymphocytes, which constitute approximately 0.01% of the total pool for B cells [68] and 0.0001% for T cells [69,70].

SARS-CoV-2 is known to spread either through respiratory or airborne routes [71]. The estimated dose transmitted via these pathways varies from several hundred to several thousand virions [72,73]. We selected a value of 1000 virions, which is notably higher than the minimum infectious dose for SARS-CoV-2 of 100 virions [74] and aligns closely with the average infectious dose.

#### 2.2.2 Time-series data

To accurately model the dynamics of viral load in the upper airways, we used time-series data provided by Sun and colleagues [75] and from a controlled study by Killingley [76], where volunteers were intranasally inoculated with SARS-CoV-2. Apart from the opportunity to investigate the infection development from the very beginning, we also got a chance to consider an incubation period, thereby significantly enhancing the biological accuracy of the model. To implement this, we added an average incubation period of 6 days [77,78] to the dynamic experimental data for all entities in the lungs, including the virus and lymphocytes. This adjustment accounts for the fact that the first and subsequent measurement points are typically recorded after the onset of symptoms, which occurs following the incubation period (Figure 2).

**Figure 2.**
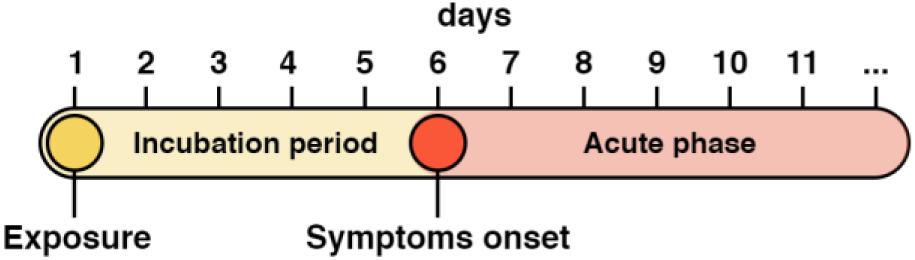
A timeline of the disease progression following exposure to the SARS-CoV-2 virus.

Based on selected experimental data, the median peak of viral load in the nose, which represents the upper airways, occurs on the 9.6^th^ day with a median of 1.1 × 10^6^ RNA copies/ml. The highest viral load was 5.86 × 10^8^and occurred on the 4^th^ day. These values are consistent with those observed by Lui and colleagues [79], who estimated the median peak to be on the 9^th^ with a median viral load of 2.5 × 10^6^RNA copies/ml. Table S3 provides additional details and other experimental data.

To model viral dynamics in the lower airways, we combined data on viral load in sputum from several studies [75,80–82]. The median peak in the combined data occurs on the 16^th^ day, with a median viral load of 2.0 × 10^6^ RNA copies/ml. The highest value, 6.84 × 10^8^, was observed on the 11^th^ day. This also corresponds to the theoretical maximum value of viral load in the lungs, estimated by Sender’s group [83].

To accurately model the antibody dynamics, we fit the model to measurements of three immunoglobulin classes (IgM, IgA, IgG) specific to the SARS-CoV-2 spike protein [84], which is known to be the main target for neutralizing antibodies [85]. Using the LOESS function to approximate the raw data, we observed that IgM peaked first on the 28^th^ day after infection, with a maximum value of 8.5 μg/ml. IgA and IgG reached their maximum levels later, on the 29^th^ day and 30^th^ day, with the highest value being approximately 10.4 μg/ml and 11.5 μg/ml for IgA and IgG respectively.

We adjusted the T cell dynamics predicted by the model to match experimental data from a retrospective study of HIV/SARS-CoV-2 coinfection, focusing only on the healthy control group, as well as data from a longitudinal study of COVID-19 patients conducted by Bergamaschi and colleagues [86,87]. According to the results, CD8+ T cells reach their peak approximately on the 18th day after infection, with a median value of 5.3 ∗ 10^4^cells/ml. Similarly, CD4+ T helper 1 type cells peak on the 23^rd^ day after exposure to the virus, with a median value of 2.8 ∗ 10^5^cells/ml. Additionally, to fit the model more accurately, we established upper boundary for biologically reliable T cell concentrations during moderate COVID-19 progression, setting it at 10^6^cells/ml for both CD4+ and CD8+ T cells [88–91].

Given the limited availability of time-series data for B cells, we established an upper boundary for the maximum biologically reliable concentration of B cells at 10^6^cells/ml [91], similar to the approach used for T cells. We suggested that the proper dynamics of B lymphocytes is indirectly ensured by fitting the model to the previously described data on immunoglobulin dynamics.

Due to the temporary nature of most cytokines, obtaining appropriate time-series data on their dynamics can be challenging. We used experimental data for IL-6, a central pro-inflammatory cytokine during COVID-19, which peaked on the 13th day after infection with a maximum value of 276 pg/ml [92]. This value also falls within the biologically plausible range, which spans from 0 to 430 pg/ml [93].

#### 2.2.3 Parameter selection

We divided the entire set of 112 parameters into 3 groups: 42 calculated parameters, 60 estimated through optimization methods, and 10 collected from other mathematical models or identified during experiments.

Among the easiest parameters to calculate are the degradation rates of cells and molecules. We used the half-life of the entity to calculate the elimination rate constant *k* (Eq. (1)), derived from the classical half-life formula [94–96].

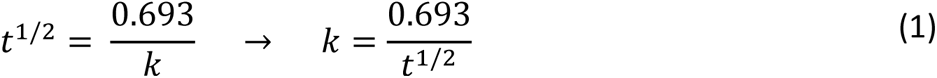

Plasma cells are known for their high productivity, with each releasing approximately 10 to 1000 antibody molecules per second [97–102]. The number of secreted antibodies depends on factors such as the immunoglobulin class, the type of plasma cell, the microenvironment, and others. For our calculations, we used immunoglobulin G as the reference molecule and assumed a fixed number of 150,000 active plasma cells per ml, which approximately corresponds to the mean number of short-lived plasma cells during an infection [91]. Considering the weight of one IgG molecule to be 150,000 Da [70,103,104], we converted the initial range of antibody secreted by plasma cell to 2.15 × 10^−7^ *to* 2.15 × 10^−5^ μg per cell per day. We then evaluated the antibody secretion rate for plasma cells per ml (1.5 × 10^5^ cells per ml), resulting in a range of 0.032 to 3.2 μg per cells/ml per day. To account for antibody dilution in the blood (5,000 ml), given that experimental data reflects serum antibody concentrations, we obtained a final range of 6.4 × 10^−6^ *to* 6.4 × 10^−4^ μg/ml per cells/ml per day. We chose the lower bound of this range to account for potential limitations in the maximum secretion rate and made minor adjustments to the final IgG secretion rate 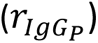 during model fitting. Values for the secretion rates 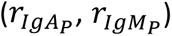 of other antibody classes were derived by varying the IgG rate within a close range.

To evaluate the rate of virion release by infected epithelial cells 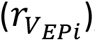, we first considered the burst size of SARS-CoV-2, defined as the number of virions produced by a single infected cell over its lifetime, ranging from 10 to 10,000 virions [83,105,106]. We then calculated the lifetime of infected cells based on their decay rate (0.5 *day*^−1^), obtaining an approximate life duration of 20 hours. This calculation yielded a range of 12 to 12,000 virions released per day. Since only about 0.01% of virions are indeed infectious [107–109], we initially opted for a rate of 1,000 virions per day and then adjusted for the infectious portion, resulting in a final rate of 10 infectious virions per day.

### 2.3 Numerical simulation

The model was solved using the Java Variable-Coefficient ODE (JVODE) solver, which is a Java implementation of the CVODE solver originally written in C [110]. JVODE is integrated into BioUML, the open-source computational platform for systems biology that we used for model development [111]. In addition to ODEs, the model incorporates delay differential equations (DDEs) to account for time delays, which are crucial for accurately modeling migration processes like the transport of dendritic cells from the lungs to the lymph nodes. To address the population heterogeneity in immune responses, we also employed stochastic differential equations (SDEs). We defined the diffusion term as a function of the variable *σ*(*t*, *x*(*t*)) = *k* ∗ *x*(*t*), where *k* is a coefficient ranging from 0 to 1, depending on the magnitude of the variable *x*(*t*). We then ran 250 simulations of the model and calculated the confidence intervals for each variable by determining the range that includes 95% of the simulated solutions. Examples of ordinary and stochastic differential equations describing virus dynamics in the upper airways are provided in Eq. (2) and (3), respectively. The stochastic part of the SDE (Eq. (3)) is represented by the diffusion term *σ*(*V*) and the Wiener process 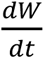.

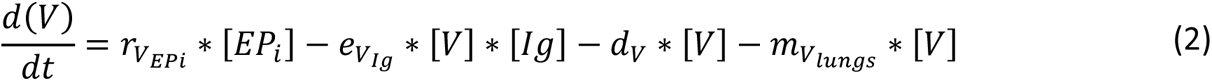

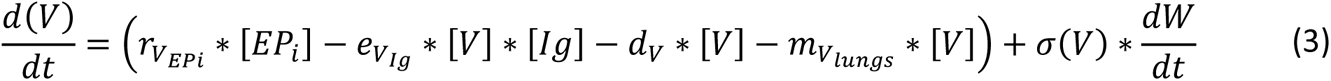

Since a substantial number of parameters could not be derived directly from experimental data or calculated, optimization methods were essential for refining the model to improve its predictive performance. We initially adjusted the model parameters manually to closely match the model dynamics with the experimental data. To identify the subset of parameters to tune, we conducted a sensitivity analysis and selected the most significant ones.

We employed three optimization algorithms implemented in the BioUML environment, including the stochastic ranking evolution strategy - SRES [112], the multi-objective particle swarm algorithm – MOPSO [113,114] and the multi-objective cellular genetic algorithm - MOCell [115,116]. Among the solutions obtained, we chose the best one (see Table S1).

### 2.4 Sensitivity analysis

To evaluate the impact of various parameters on the progression of COVID-19, we performed a local sensitivity analysis. We used the cumulative viral load throughout the course of infection as a measure of COVID-19 severity:

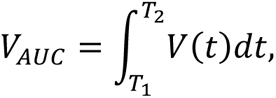

where T_1_ represents the time of virus appearance and T_2_ represents the time of complete virus clearance.

To calculate the local sensitivity indices, we computed partial derivatives with respect to the vector of model parameters using the finite difference approximation method, which is implemented in BioUML environment:

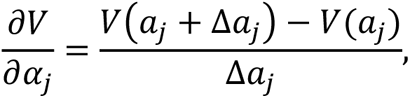

where Δ*a*_*j*_ represents a small perturbation to the local parameter *a*_*j*_, *V*(*a*_*j*_ + Δ*a*_*j*_) and *V*(*a*_*j*_) correspond to the solutions of the algebraic systems *f*(*V*, *a*_*j*_ + Δ*a*_*j*_) = 0 and *f*(*V*, *a*_*j*_) = 0, respectively.

Due to the variability in model parameters spanned eighteen orders of magnitude, we normalized the obtained values by multiplying them by a normalization factor 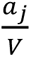. The sensitivity analysis was conducted on a model fitted to moderate COVID-19 progression.

## 3. Results

### 3.1 Local sensitivity analysis

To assess the model’s response to parameter perturbations, we conducted a local sensitivity analysis and categorized the parameters with the most significant impact on the simulation outcome into four groups based on their biological roles: infection, innate immunity, adaptive immunity, and cytokines (Figure 3). We found that the cumulative viral load, which represents the total amount of virus over the course of infection (i.e., the area under the viral load curve), positively correlates with parameters primarily associated with the virus and the infection process. These includes the rate of virion release by infected epithelial cells 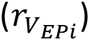 in both the upper airways and lungs, the rate of epithelial cell infection by the virus 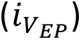, and the parameter 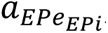, which is related to the incubation period of infected epithelial cells. On the other hand, a negative correlation was observed with parameters associated with the immune response. The most significant parameters are related to adaptive immunity, including the rate of naïve CD4+ T cell proliferation (*p*_*CD*4_), the elimination of infected cells by CTLs 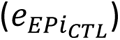, and the migration rate of CTLs to the lungs 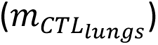. Another crucial parameter pertains to the maturation and migration of immature dendritic cells to the lymph nodes 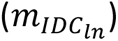, emphasizing the role of innate immunity. Collectively, these findings highlight the importance of both innate and adaptive components of the immune response in regulating the progression of the infection.

**Figure 3.**
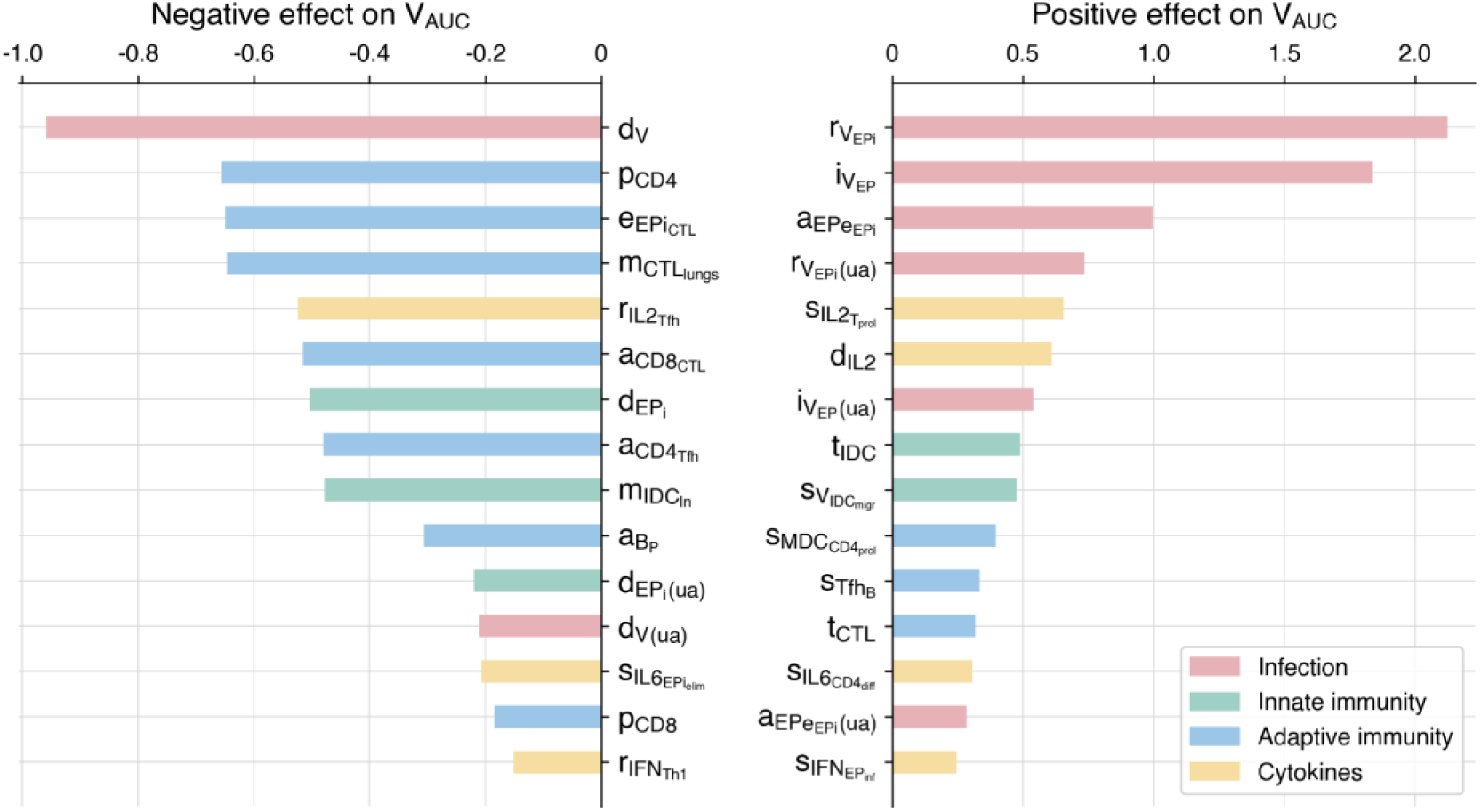
Scaled sensitivity coefficients that have negative (right) and positive (left) effects on the cumulative viral load.

Additionally, we assessed the sensitivity coefficients for epsilon parameters, which determine the efficacy of age-related processes (Figure 4). Details on the epsilon parameters and their significance are provided in Section 3.3. As expected, a decrease in the values of epsilon parameters, reflecting aging, results in increased viral load. The most significant parameters are related to the proliferation of naïve CD4+ T cells (*ɛ*_1_), the elimination of infected epithelial cells by CTLs (*ɛ*_7_), the differentiation of naïve CD8+ T cells to CTLs (*ɛ*_3_), and the activation, migration, and maturation of immature dendritic cells (*ɛ*_5_).

**Figure 4.**
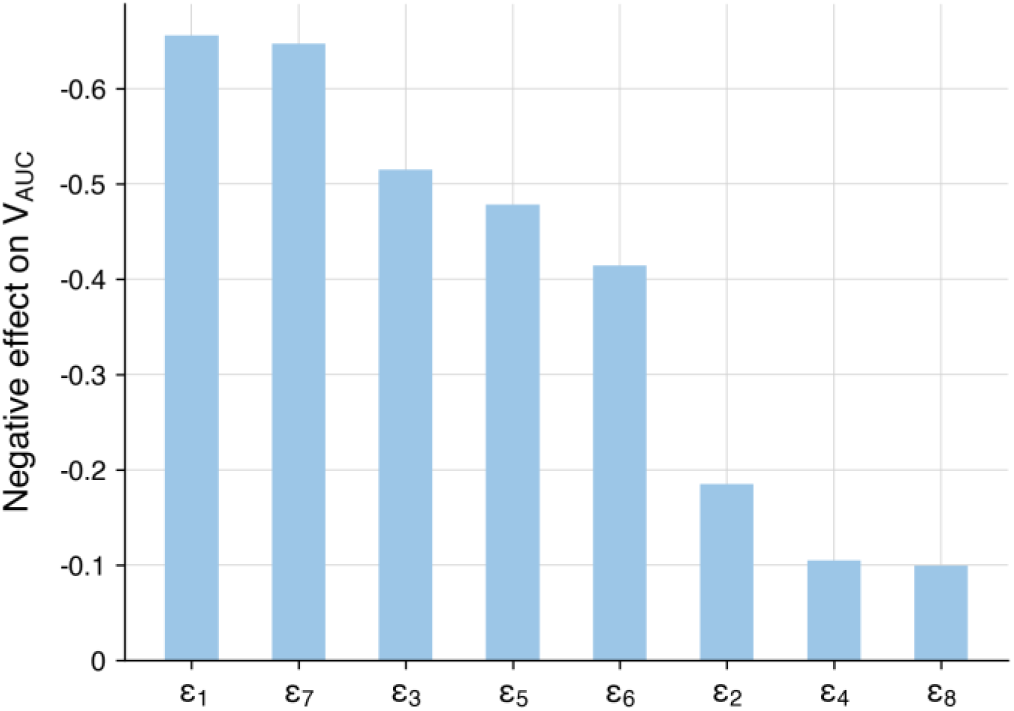
Scaled sensitivity coefficients for **ε** parameters for cumulative viral load.

### 3.2 Baseline model and optimization

We initially fitted the developed modular model to experimental data of moderate disease progression. Details on the parameters and initial values for this moderate version are provided in Section 2.2, Table S1, and Table S2. This model was used as the baseline for further analyses and subsequent modifications.

Solutions from the fitted baseline model align well with the experimental data (Figure 5). The peak of viral load in the upper airways and lungs falls on the 9^th^ and 12^th^ days, with values of 3.76 × 10^7^ and 9.42 × 10^7^ RNA copies/ml respectively. These results not only match clinical data but also correspond to the theoretically calculated values by Sender and colleagues [83]. The complete elimination of the virus from the body occurs by the 24^th^ day after infection, which corresponds to the median SARS-CoV-2 clearance by 22^nd^ day after infection [116].

**Figure 5.**
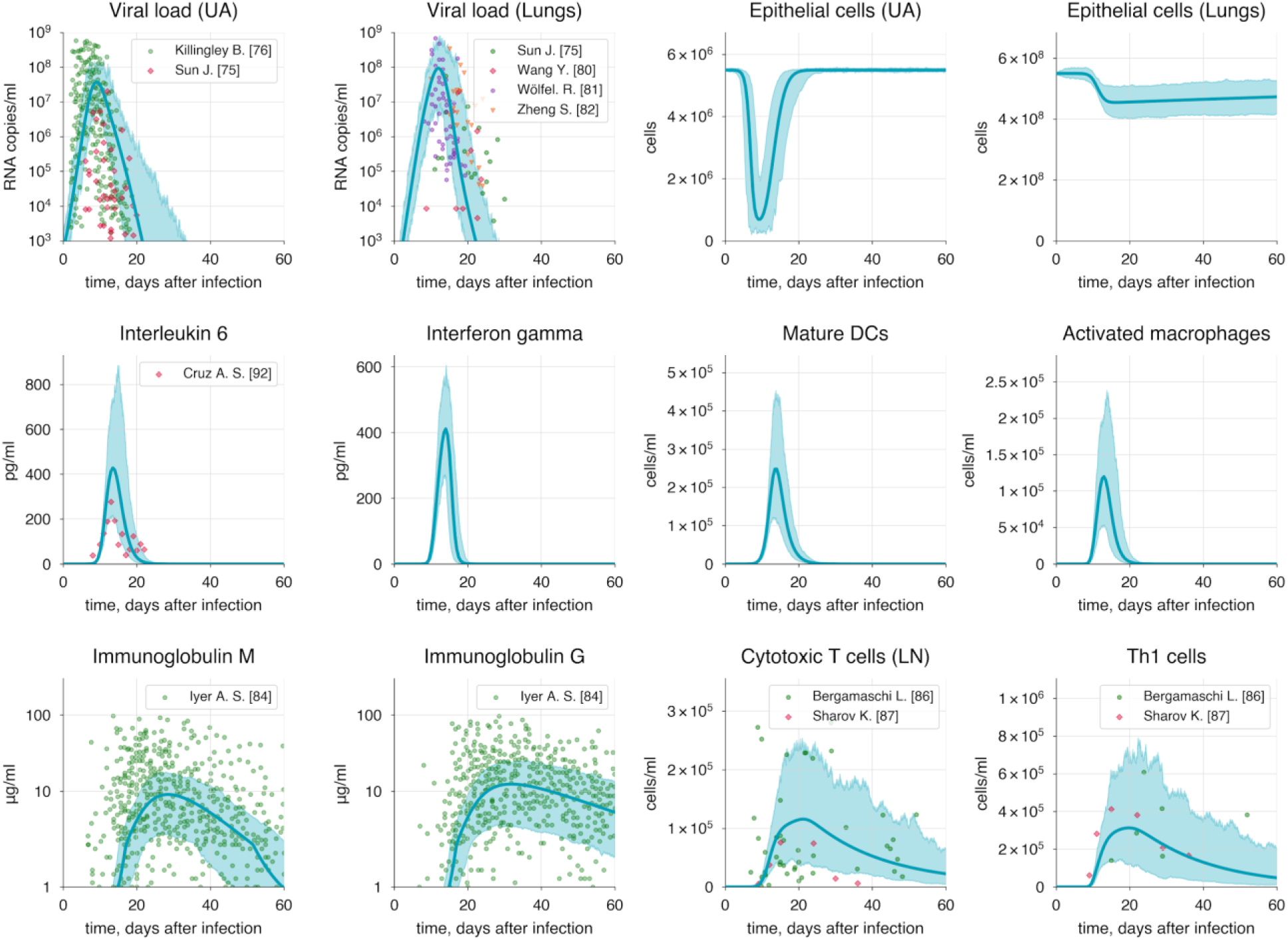
Baseline solution of the model. Lung variables are plotted, except for epithelial cells and viral load, which are shown for both compartments: upper airways (UA) and lungs. Extended versions of all subsequent plots are given in S1 Appendix (Figures 1-2). The blue shaded area represents the 95% confidence interval.

Viral shedding is known to begin about 2 days before symptom onset [118,119], which corresponds to the 4^th^ day after infection in our model. We used the viral load in the upper airways on the 4^th^ day (10^5^RNA copies/ml) as an approximate threshold for the onset and end of shedding (Figure 6). Consequently, we determined the viral shedding period to be from 4 to 17 days after infection, resulting in a total duration of 13 days, which is consistent with the median viral shedding duration of 11 days for moderate COVID-19 [120].

**Figure 6.**
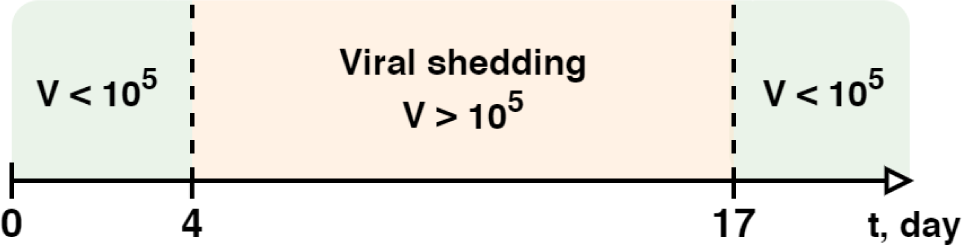
Viral shedding duration in the baseline model.

Since lung damage is a key indicator of the disease severity, we also tracked the numbers of healthy and infected epithelial cells. At the peak of the infection, the number of infected cells reaches a maximum of 8.9 × 10^6^cells, corresponding to the theoretical peak calculated by Sender and co-authors [83]. The total alveolar epithelial damage reaches 17.3%, corresponding to 4.55 × 10^8^. These results are consistent with the criteria for moderate COVID-19 progression (see Table 2).

The major components of the humoral immune response, IgM, IgA and IgG, peaked on the 28^th^, 29^th^ and 31^th^ day after infection, with the highest values reaching 9.0 μg/ml, 11.0 μg/ml and 12.4 μg/ml, respectively. Additionally, the seroreversion time – the period required for antibodies in the serum to completely disappear or decrease to undetectable levels – was consistent with clinical data, being 60, 72 and 102 days for IgM, IgA and IgG correspondingly. The growth of antibodies is driven by the development of plasma cells, which peak on the 22^nd^ day after infection.

CD4+ T cells peaked on the 20^th^ and 22^nd^ day, reaching 3.9 × 10^5^ cells/ml and 2.2 × 10^5^ cells/ml for Th1 and Tfh respectively. In contrast, CTLs peaked on the 21^st^ day with a maximum value of 1.2 × 10^5^ cells/ml. The median values for Th1 cells (1.6 × 10^5^ cells/ml) and CTLs (6.5 × 10^4^cells/ml), along with the median peak times (20^th^ and 21^th^ days, respectively), closely align well with the experimental data discussed in Section 2.2.

The IL-6 levels we observed align closely with the experimental data, peaking on the 13^th^ day at a maximum of 430 pg/ml. Other cytokines, IL-2, IL-12, and IFNγ peaking on the 20^th^, 13^th^, and 14^th^ days with concentrations of 10 pg/ml, 70 pg/ml, and 400 pg/ml, respectively.

Macrophages and dendritic cells begin to activate almost immediately after the virus becomes detectable, bridging the innate and adaptive immunity. Activated macrophages peak on the 13^th^ day with a value of 1.2 × 10^5^cells/ml, while mature dendritic cells reach their peak on the 14^th^ day at 2.5 × 10^5^ cells/ml.

### 3.3 Modes of the immune response during COVID-19

The progression of COVID-19 is significantly influenced by risk factors such as age, obesity, or diabetes [121]. Among these, age is shown to be a key predictor of severe or critical COVID-19 outcomes due to gradual changes in the immune system associated with aging [122,123]. Immunosenescence, the aging of the immune system, begins around age 20 and progresses becoming significantly noticeable by age 60 [123]. Senescence affects various aspects of the immune system with a predominant impact on adaptive immunity. It has been demonstrated that in elderly individuals the stimulation of naïve B cells by dendritic cells is reduced by up to 70%, which impairs B cell proliferation and, consequently, the formation of antibodies [124,125]. This is further complicated by age-related impairments in germinal centers (GCs) response, which involve reduction in both size and function of GCs [126]. Similar changes affect naïve T cells leading to diminished expansion and differentiation capabilities [127–130]. Additionally, the ability of cytotoxic T cells to eliminate defective cells also declines with age [131], thereby reducing the overall efficiency of the immune response and contributing to a longer and more severe disease progression. Moreover, studies on animal models show that antibodies produced by aged mice not only have lower affinity and avidity compared to those from younger mice but also exhibit reduced protective efficacy against pathogens [132]. These findings also have been confirmed in humans [8].

However, a dramatic decline in the number of naïve T and B cells remains a primary age-related impairment of the immune system [134,135]. This significantly affects the body’s protective abilities due to a reduced number of antigen-specific immune cells, which are essential for mounting a robust immune response. Furthermore, immunosenescence affects not only adaptive immunity but also innate immunity. Research suggests that dendritic cells and macrophages have a diminished ability to be activated by processing antigens and to stimulate other immune cells. Additionally, impaired expression of homing factors in lymph nodes, which are essential for the transfer of antigen-presenting cells to lymph nodes, significantly reduces the migration potential of dendritic cells [127,135].

Following the National Institutes of Health guidelines [3], we categorized potential COVID-19 progressions into three modes: moderate, severe, and critical. We use the moderate progression as the baseline model, with detailed information provided in Section 3.2. Since the severity of COVID-19 is heavily influenced by risk factors mentioned earlier, we incorporated all identified age-related immune system alterations into a parameter vector *ɛ* = (*ɛ*_1_, *ɛ*_2_, *ɛ*_3_, *ɛ*_4_, *ɛ*_5_, *ɛ*_6_, *ɛ*_7_, *ɛ*_8_) defining the effectiveness of these processes. Each parameter initially has a value of 1, representing no change, which is standard for the baseline model. Detailed description of the processes associated with each parameter is provided in Table 1. By decreasing these parameters from 1 (baseline) to 0 and lowering the initial populations of naive T and B cells in both the lungs and upper airways, we simulated various disease progression scenarios (Table 3).

**Table 1.** Age-related processes and associated parameters.

| Parameter | Description of the process | References |
| --- | --- | --- |
| $\varepsilon_1$ | Proliferation of naïve CD4+ T cells | [127–129,136] |
| $\varepsilon_2$ | Proliferation of naïve CD8+ T cells | |
| $\varepsilon_3$ | Naïve CD8+ T cell differentiation into CTL | |
| $\varepsilon_4$ | Proliferation of naïve B cells | [124,125] |
| $\varepsilon_5$ | Immature dendritic cell maturation and migration to the lymph nodes | [127] |
| $\varepsilon_6$ | Virion neutralization by immunoglobulins (IgA, IgG, IgM) | [132] |
| $\varepsilon_7$ | Infected epithelial cell elimination by CTLs | [131] |
| $\varepsilon_8$ | Activation of resting macrophages | [135] |

**Table 2.**
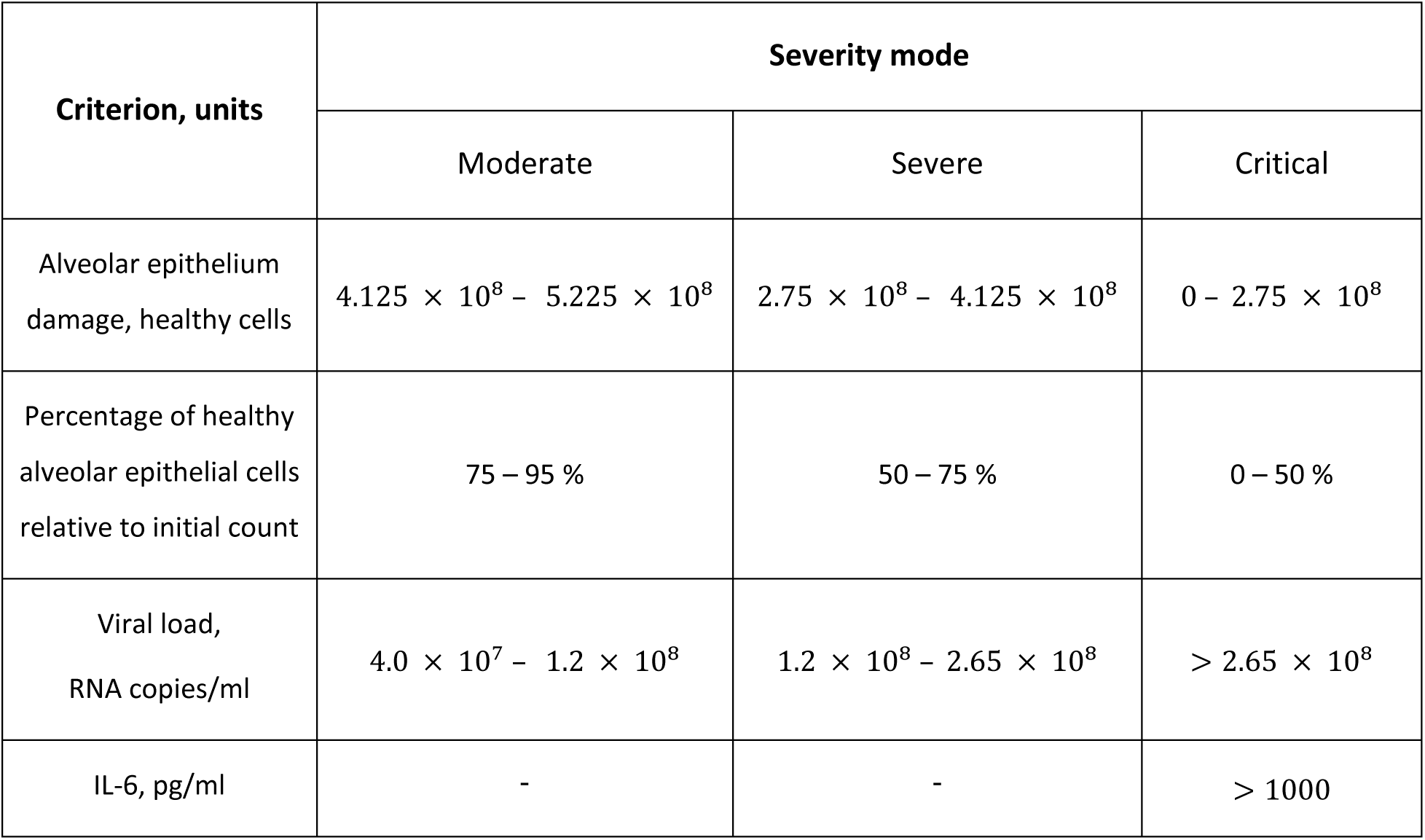
Criteria for disease severity modes.

| Criterion, units | Severity mode |  |  |
| --- | --- | --- | --- |
|  | Moderate | Severe | Critical |
| Alveolar epithelium damage, healthy cells | $4.125 \times 10^8 - 5.225 \times 10^8$ | $2.75 \times 10^8 - 4.125 \times 10^8$ | $0 - 2.75 \times 10^8$ |
| Percentage of healthy alveolar epithelial cells relative to initial count | 75 – 95 % | 50 – 75 % | 0 – 50 % |
| Viral load, RNA copies/ml | $4.0 \times 10^7 - 1.2 \times 10^8$ | $1.2 \times 10^8 - 2.65 \times 10^8$ | $> 2.65 \times 10^8$ |
| IL-6, pg/ml | - | - | > 1000 |

**Table 3.** Parameter values for disease severity modes.

| Parameter | Severity mode |  |  |
| --- | --- | --- | --- |
|  | Moderate | Severe | Critical |
| $\epsilon_1$ | 1 (100%) | 0.9 (90%) | 0.8 (80%) |
| $\epsilon_2$ | 1 (100%) | 0.9 (90%) | 0.8 (80%) |
| $\epsilon_3$ | 1 (100%) | 0.9 (90%) | 0.8 (80%) |
| $\epsilon_4$ | 1 (100%) | 0.9 (90%) | 0.8 (80%) |
| $\epsilon_5$ | 1 (100%) | 0.9 (90%) | 0.8 (80%) |
| $\epsilon_6$ | 1 (100%) | 0.9 (90%) | 0.8 (80%) |
| $\epsilon_7$ | 1 (100%) | 0.9 (90%) | 0.8 (80%) |
| $\epsilon_8$ | 1 (100%) | 0.9 (90%) | 0.8 (80%) |
| $B_n$ | 60000 (100%) | 48000 (80%) | 42000 (70%) |
| $T_n$ | 33000 (100%) | 26400 (80%) | 23100 (70%) |
| $H_n$ | 100000 (100%) | 80000 (80%) | 70000 (70%) |
| $B_n(\text{ua})$ | 16000 (100%) | 12800 (80%) | 11200 (70%) |
| $T_n(\text{ua})$ | 33000 (100%) | 26400 (80%) | 23100 (70%) |

We selected specific indicators to distinguish between different severities of COVID-19, including the number of damaged epithelial cells, viral load, and interleukin-6 concentration. This choice is supported by numerous studies [137–149] demonstrating a correlation between these markers and disease severity that is discussed below. The simulation of different COVID-19 scenarios is presented in Figure 7.

**Figure 7.**
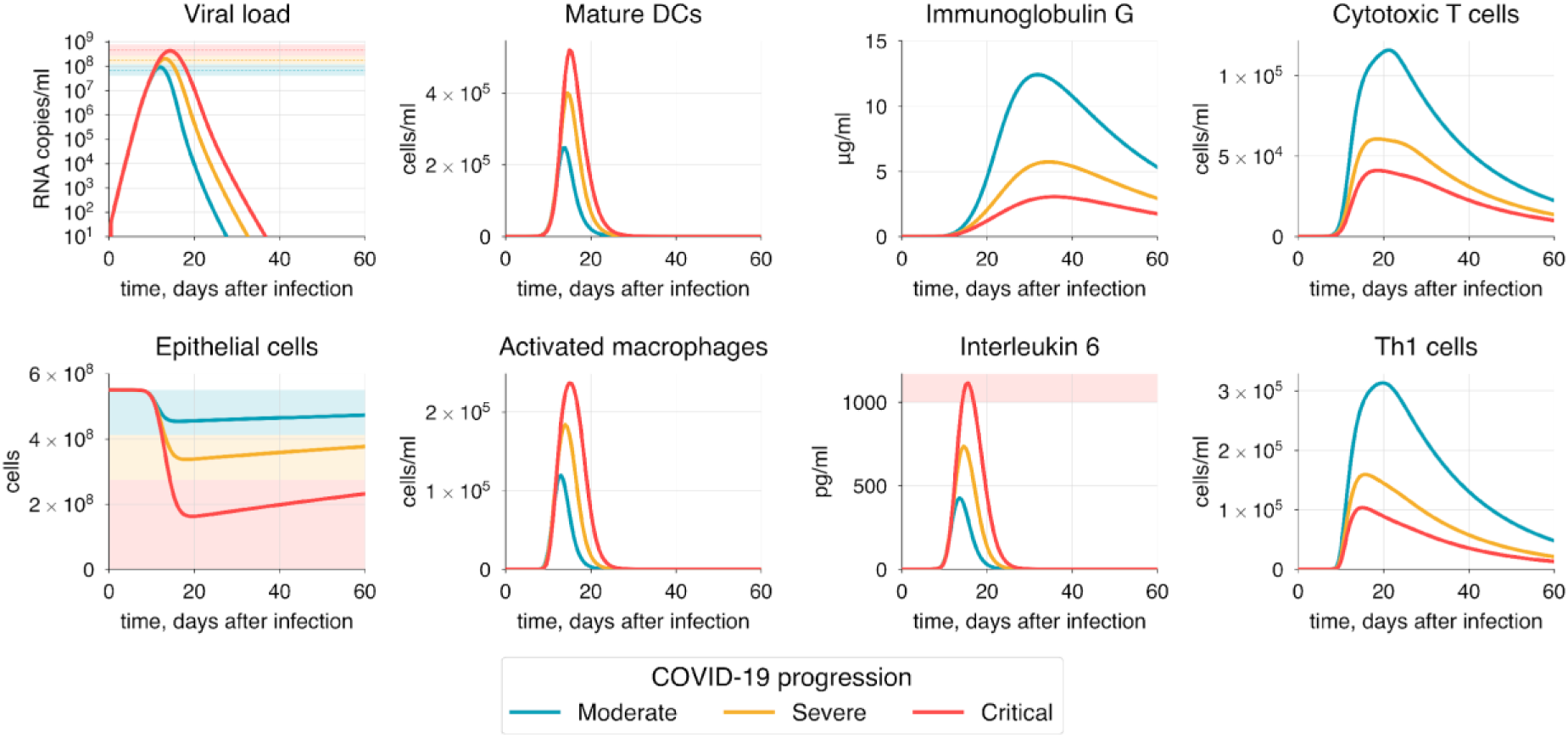
The simulation of moderate, severe, and critical COVID-19 progressions. Shaded areas in some plots indicate severity levels: green for moderate, yellow for severe, and red for critical progression.

Diffuse alveolar damage (DAD) is a frequent histological hallmark of COVID-19, often necessitating mechanical ventilation and potentially leading to death [137]. This occurs because DAD disrupts normal gas exchange within the alveoli, triggering inflammation and subsequent fibrosis [138]. Post-mortem studies suggest a high probability of fatal outcomes when lung involvement exceeds 50% [139,140]. Therefore, to differentiate disease severity, we established lung damage thresholds as detailed in Table 2.

Studies have shown that viral load is significantly higher in patients with severe and critical conditions compared to those with moderate disease [141,142]. Severe patients typically have a viral load about 2-3 times greater than that of moderate cases [143,144]. We defined a median viral load for moderate progression as 7.0 × 10^7^ RNA copies/ml [145]. Based on this, we propose median viral loads for severe and critical cases of approximately 1.8 × 10^8^ RNA copies/ml and 4.6 × 10^8^ RNA copies/ml, respectively. Additionally, we consider viral shedding in relation to disease severity, as evidence suggests that patients with more severe symptoms may experience a longer duration of viral shedding compared to those with milder COVID-19 [146–148], as discussed in Section 3.2.

One of the most significant hallmarks of severe COVID-19 is interleukin-6, which has been shown to strongly correlate with disease severity [148]. In most cases, IL-6 levels exceeding 1000 pg/ml are associated with a higher risk of death [149].

To achieve the specific COVID-19 mode, we adjusted the vector of parameters ε, as previously discussed, focusing on the criteria established for each severity state. It is important to note that achieving a particular disease progression can occur through various combinations of epsilon parameter values and the initial number of naïve T and B cells. This variability reflects the heterogeneity in the immunosenescence process among individuals in the population. Here, we propose one option for each disease severity state.

### 3.4 Validation of the model

To evaluate the effectiveness of our model in predicting COVID-19 outcomes, we explored how various aspects of the immune response and mechanisms of the infection influence COVID-19 progression and severity. Furthermore, we examined different treatment strategies, including those aimed at enhancing the immune response and inhibiting viral replication. We also addressed the impact of immunosuppression in individuals who have undergone transplantation and those with immunosuppressive conditions such as HIV. All conducted experiments are presented in Table 4.

**Table 4.**
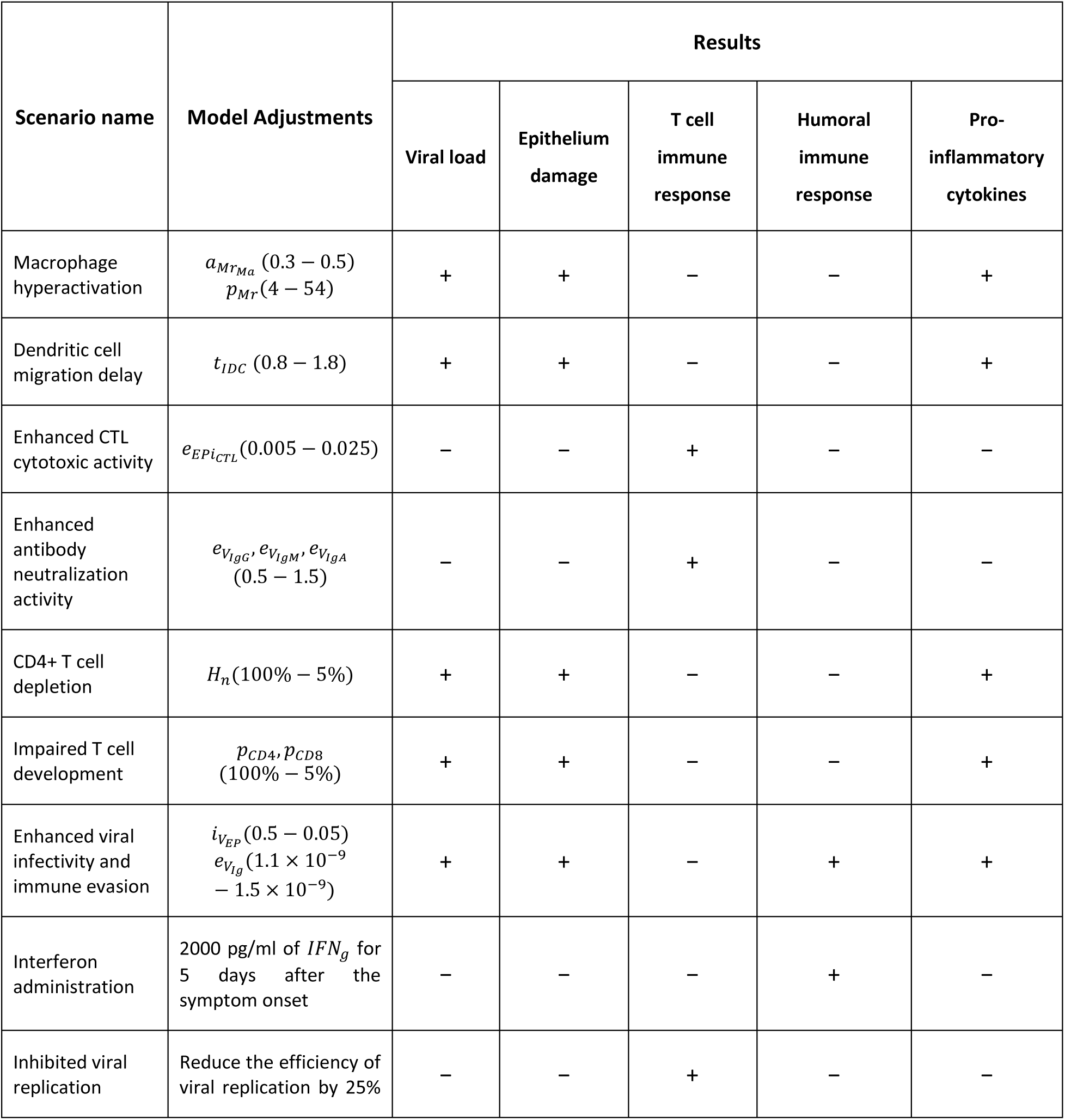
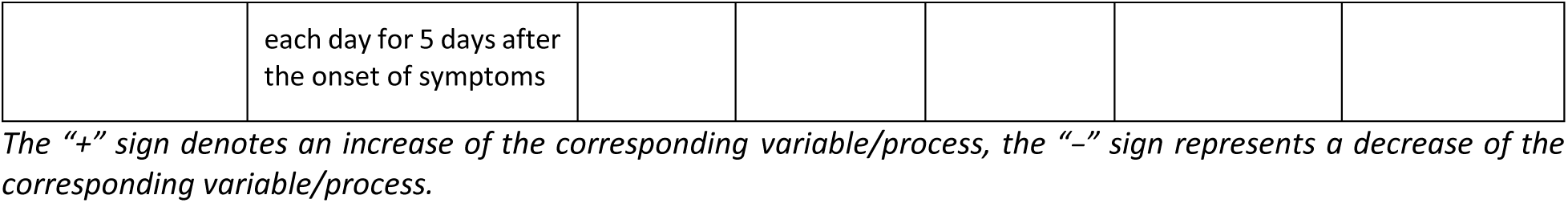
Validation scenarios.

#### 3.4.1 Immune response

One of the most significant hallmarks of severe COVID-19 is the hyperactivation of the innate immune response, which leads to a substantial increase in the release of pro-inflammatory cytokines [150,151]. This results in prolonged inflammation and suppression of the adaptive immunity, particularly affecting T cells. Interleukin-6 (IL-6) is a key player in this process [148,152], as evidenced by the correlation between IL-6 levels and disease severity. To simulate this, we gradually increased the rates of macrophage activation 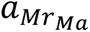, and macrophage recruitment *p*_*Mr*_(Figure 8).

**Figure 8.**
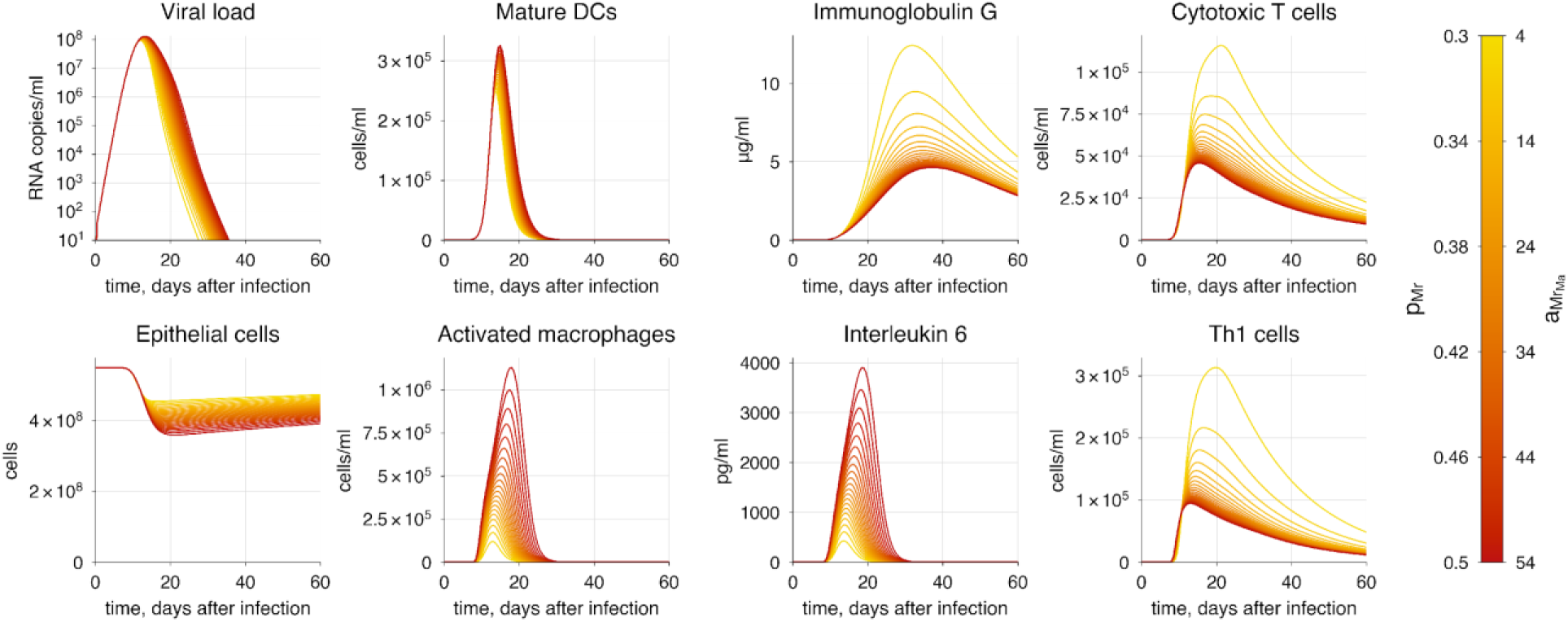
The baseline model solution with macrophage hyperactivation. The varying parameters represent the rates of macrophage recruitment and activation. The transition from yellow to red indicates increasing severity of COVID-19.

We observed an increase in viral load and duration of viral shedding, indicating a worse disease outcome. We also noted that CTL and antibody responses are impaired by significantly elevated levels of IL-6. Thus, these findings suggest that macrophage hyperactivation is directly proportional to the severity of COVID-19.

In addition to hyperactivation of the innate immune response, a study revealed a strong correlation between COVID-19 severity and delay in the innate immune response [153]. At *in silico* experiment we increased the delay of IDC maturation and migration to the lymph nodes (*t*_*IDC*_) for up to 24 hours. We observed an increased viral load and greater alveolar epithelium damage (Figure 9). Additionally, we noted a corresponding shift in the peaks of T cell and antibody responses, which allows the virus to reproduce more freely during the initial phase of the infection.

**Figure 9.**
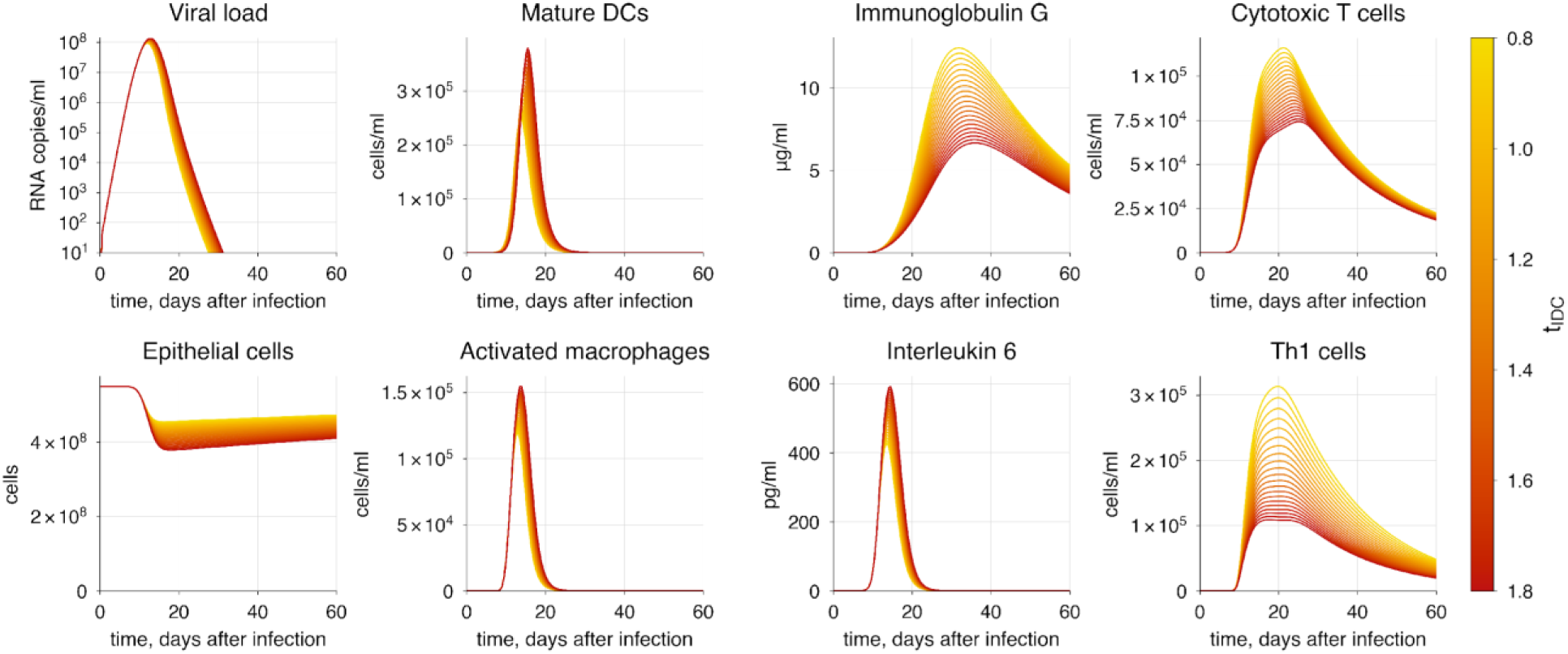
The baseline model solution with innate immune response delay. The varying parameter represents the delay of dendritic cell maturation and migration to lymph nodes. The transition from yellow to red indicates increasing severity of COVID-19.

Furthermore, we investigated how viral load depends on the elimination rate of infected epithelial cells by cytotoxic T cells 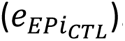. The solution demonstrates a decrease in virus concentration and epithelial cell damage (Figure 10), leading to a better outcome and results in a milder course of the disease, accompanied by a reduction in the intensity of the immune response.

**Figure 10.**
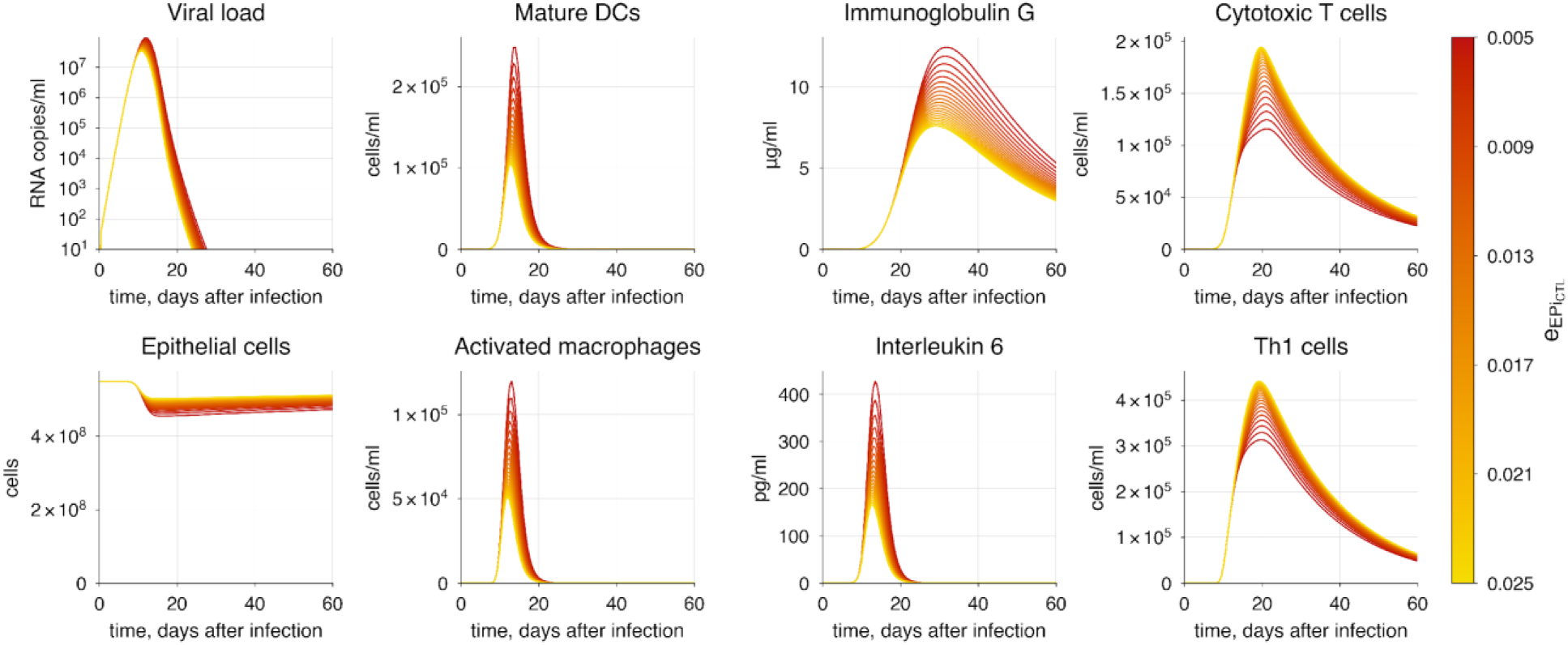
The baseline model solution with enhanced CTL-mediated infected cell clearance. The varying parameter represents the rate of infected cell elimination by CTLs. The transition from red to yellow indicates decreasing severity of COVID-19.

A similar pattern to that observed with T cells was noted during the increase in antibody neutralization activity, though changes in viral load was less significant (Figure 11). This is expected, as antibodies typically peak closer to the point of viral clearance, while T cells primarily suppress the infection during the acute phase. In contrast, antibodies provide immunity throughout their entire period of seroreversion. At the experiment, we simultaneously varied the neutralization constants for all three antibody classes (IgM, IgA, IgG).

**Figure 11.**
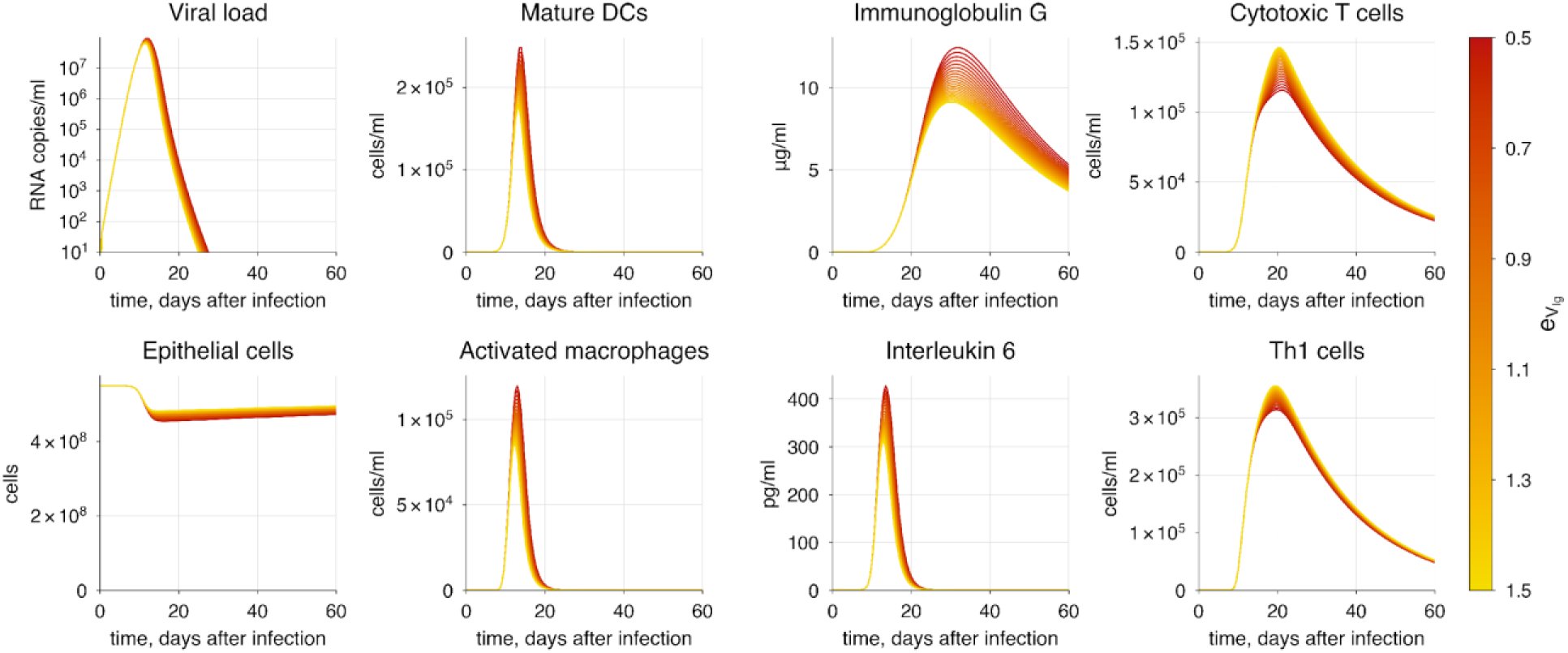
The baseline model solution with enhanced antibody-mediated virion neutralization. The varying parameter represents the rate of virion neutralization by antibodies. The transition from red to yellow indicates decreasing severity of COVID-19.

#### 3.4.2 Immunosuppression

We identified and examined two types of immunosuppression: one resulting from HIV infection and the other from immunosuppressive therapy, employed in organ transplantation and autoimmune diseases [154,155]. The first type involves the depletion of CD4+ T cells [156], weakening the adaptive immune response and making the body dramatically vulnerable to infectious agents like SARS-CoV-2. At the second case, immunosuppression is achieved through medication primarily aimed at reducing T cell proliferation, resulting in substantially lower T cell concentrations, sometimes leading to their complete depletion [157]. Studies indicate a significantly higher risk of poor COVID-19 outcomes in patients with solid organ transplantation [43,159], placing immunosuppressed individuals in a high-risk group.

To simulate the first scenario, we gradually decreased the initial concentration of naïve CD4+ T cells from 100% to 5% of the initial value. This reflects the progression of HIV infection, ultimately leading to the AIDS state characterized by the complete depletion of CD4+ T cells [159]. The simulation demonstrated a predictable increase in viral load and epithelium damage, resulting from the development CD4+ T cell lymphopenia, which affects both cellular and humoral immune responses (Figure 12). This corresponds to the research findings reporting impaired plasma cell formation and exhaustion of CD8+ T cells, which, combined with SARS-CoV-2 infection, contribute to critical conditions where the emerging immune response is insufficient to combat the virus [161–163]. According to the criteria for critical lung tissue damage incompatible with life (Table 2), we conclude that HIV positive patients with a reduction CD4+ T cell counts by more than 50% from the initial level are unlikely to survive COVID-19.

**Figure 12.**
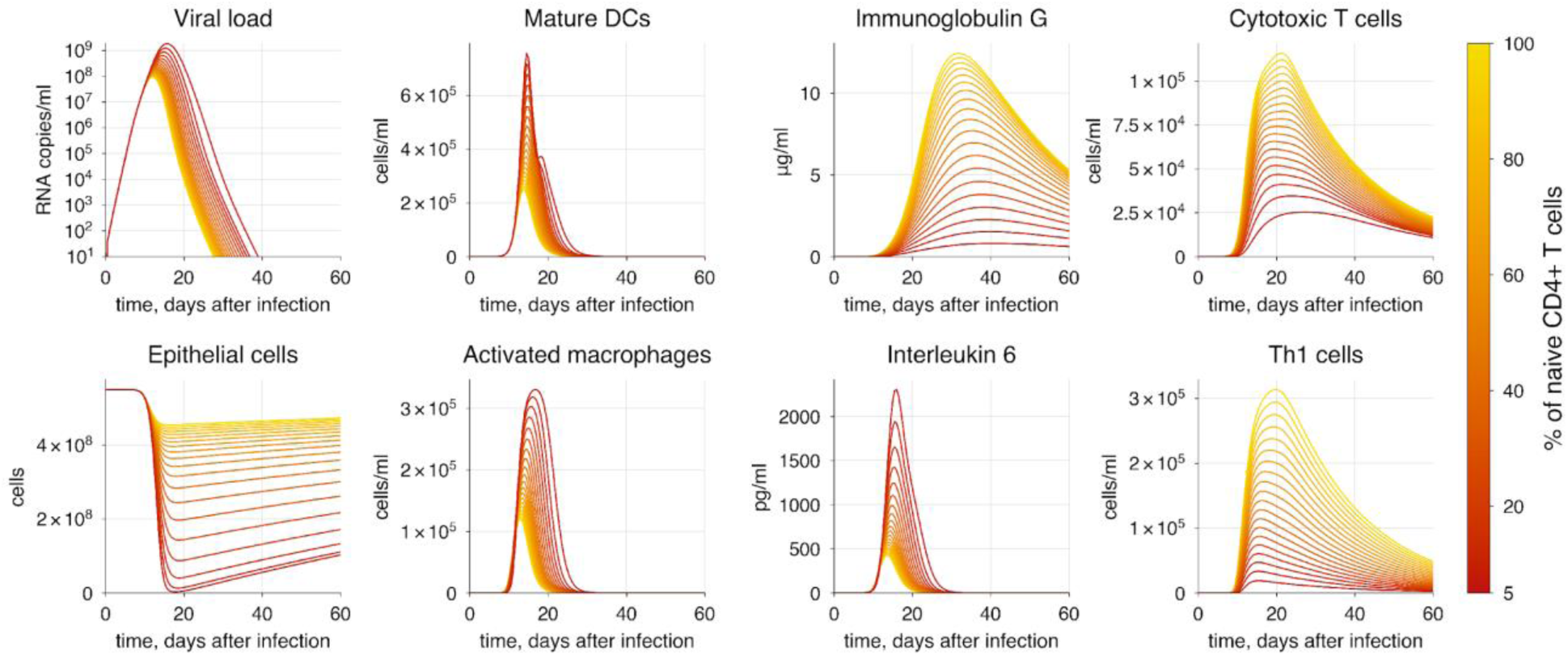
The baseline model solution with naive CD4+ T cell depletion. The transition from yellow to red indicates increasing severity of COVID-19.

To simulate the second scenario, we chose corticosteroids as an example of a drug representing a classical strategy for immunosuppression therapy. Corticosteroids significantly impact adaptive immunity, particularly affecting T cell development [164,165], while they do not substantially decrease B cells [165]. Thus, we gradually reduced the proliferation rates of both CD4+ (*p*_*CD*4_) and CD8+ (*p*_*CD*8_) T cells.

A significant reduction in T cell proliferation rates results in the inability of the immune system to completely eliminate the virus from the body (Figure 13). When T cell proliferation efficiency falls below 40% of initial values, it leads to virus persistence and a dramatic decline in the number of alveolar cells as the immune response wanes. At the same time, we observed a predictable worsening of the disease outcome, which is directly proportional to the extent of T cell proliferation impairment.

**Figure 13.**
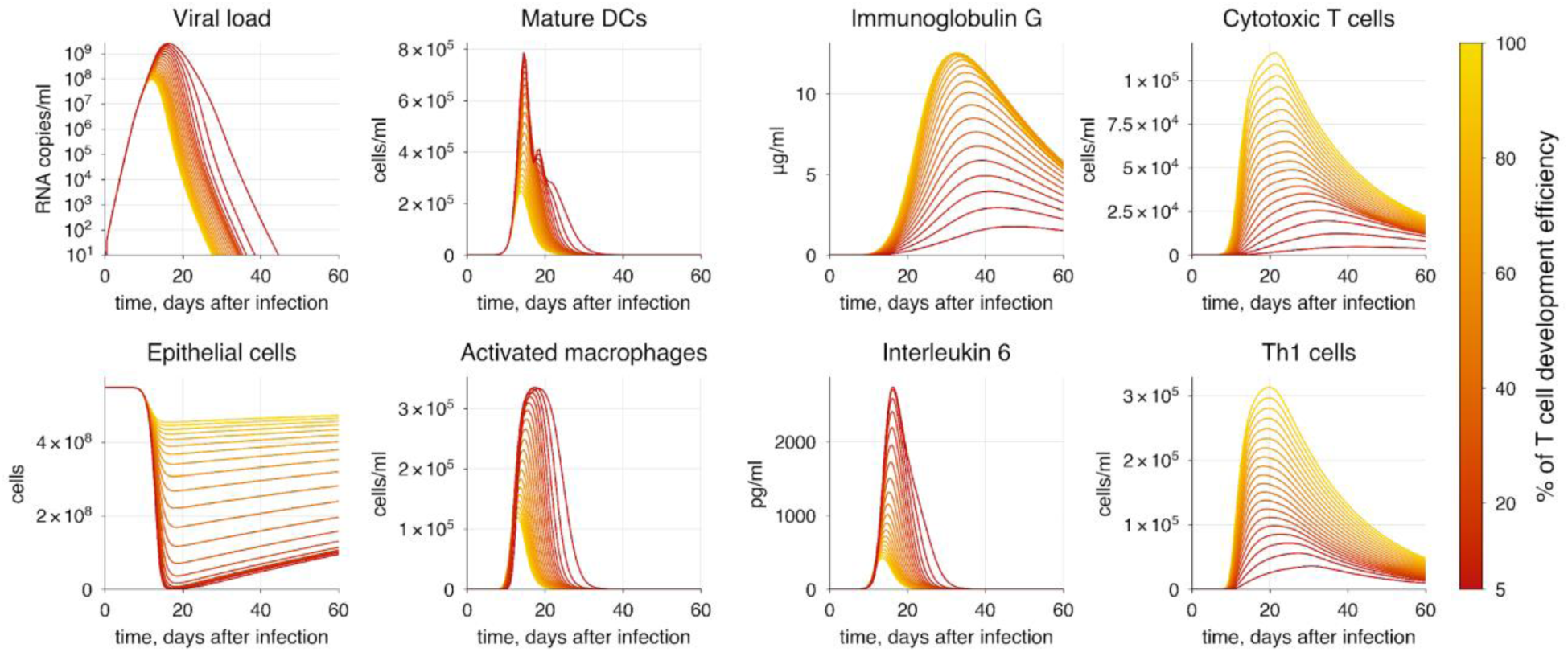
The baseline model solution with impaired T cell development due to immunosuppression. The varying parameters represent the rates of CD4+ and CD8+ T cell proliferation. The transition from yellow to red indicates increasing severity of COVID-19.

Thus, this raises important issues regarding the need for a more careful approach to immunosuppressive cases and underscores the importance of developing and implementing new treatment methods.

#### 3.4.3 SARS-CoV-2 infectivity

Since the onset of the pandemic many variants of SARS-CoV-2 have emerged. These new viral strains commonly exhibit increased transmissibility and lead to more severe disease compared to the wild type. Beyond that, the virus has shown the ability to evade human immunity, particularly targeting the humoral immune response, resulting in a reduction of antibody neutralization activity by more than 50% in some cases. These characteristics seem to be valid for all previously dominant SARS-CoV-2 variants: Alpha, Beta, Gamma and Delta [167–170], but have changed in more recent variants such as Omicron. In contrast, it reliably causes less severe symptoms [170,171] while exhibiting significantly higher contagiousness [172], which altogether allowed it to become the dominant SARS-CoV-2 variant.

Since the experimental data we used are not associated with a specific viral strain, we focused on investigating the effects of varying parameters related to cell infection and antibody evasion, treating these parameters as factors determining the viral strain.

We examined the concurrent increase in both the rate of epithelial cell infection 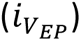 and virion evasion from neutralizing antibodies 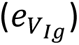. During the experiment we observed an increase in viral load and damage of alveolar epithelium (Figure 14). Moreover, the duration of viral shedding expanded as well, indicating not only a more severe progression of COVID-19 but also enhanced virus transmissibility.

**Figure 14.**
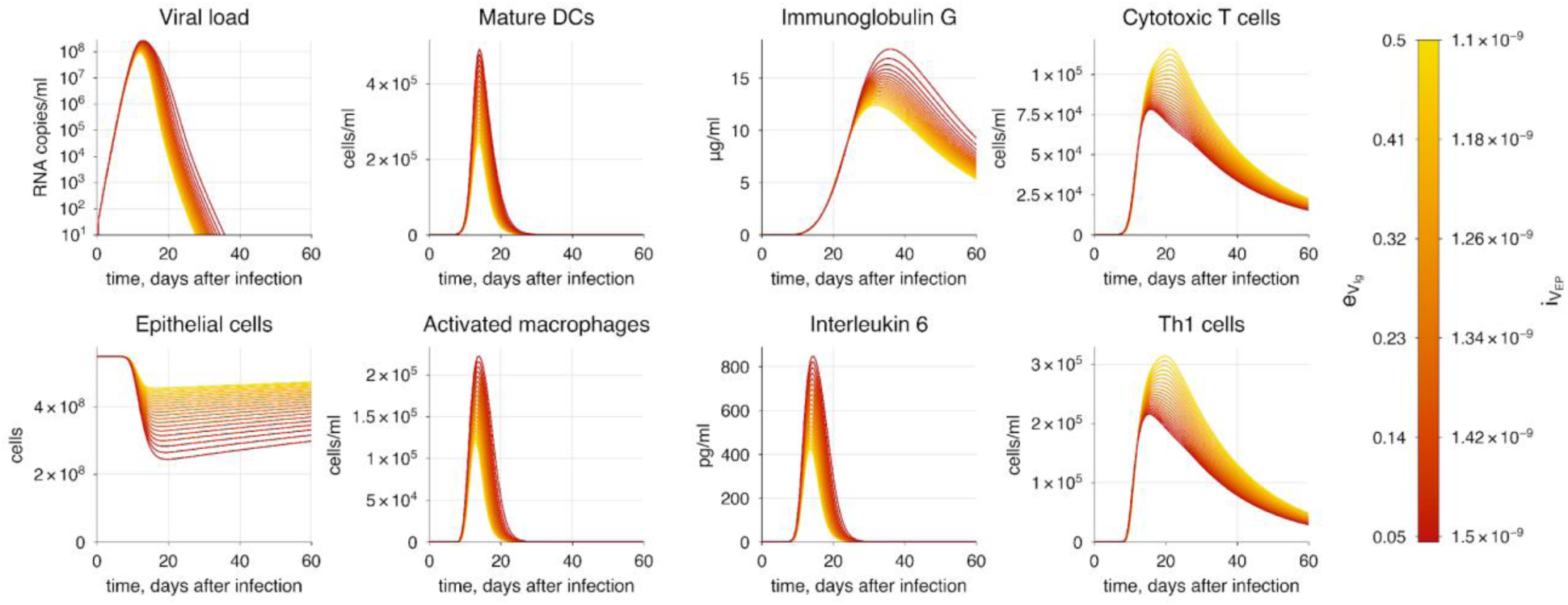
The baseline model solution with enhanced viral infectivity and immune evasion. The varying parameters represent the rates of epithelial cell infection by the virus and virion neutralization by antibodies. The transition from yellow to red indicates increasing severity of COVID-19.

#### 3.4.4 Treatment strategies

Interferons are well-known for their role as key inducers of intracellular immunity, helping cells combat pathogens before the activation of the primary immune response [173–175]. We implemented this mechanism through IFN-dependent inhibition of epithelial cell infection (see S1 Text). To study the dynamic changes in immune response resulting from IFN action, we followed a treatment protocol [176] in which patients received daily doses of interferon at a concentration of 2000 pg/ml for five days, starting immediately after the symptom onset. Compared to patients who did not receive any treatment, those who did experienced more than a 5-fold reduction in viral load and a predictable decrease in the disease severity (Figure 15).

**Figure 15.**
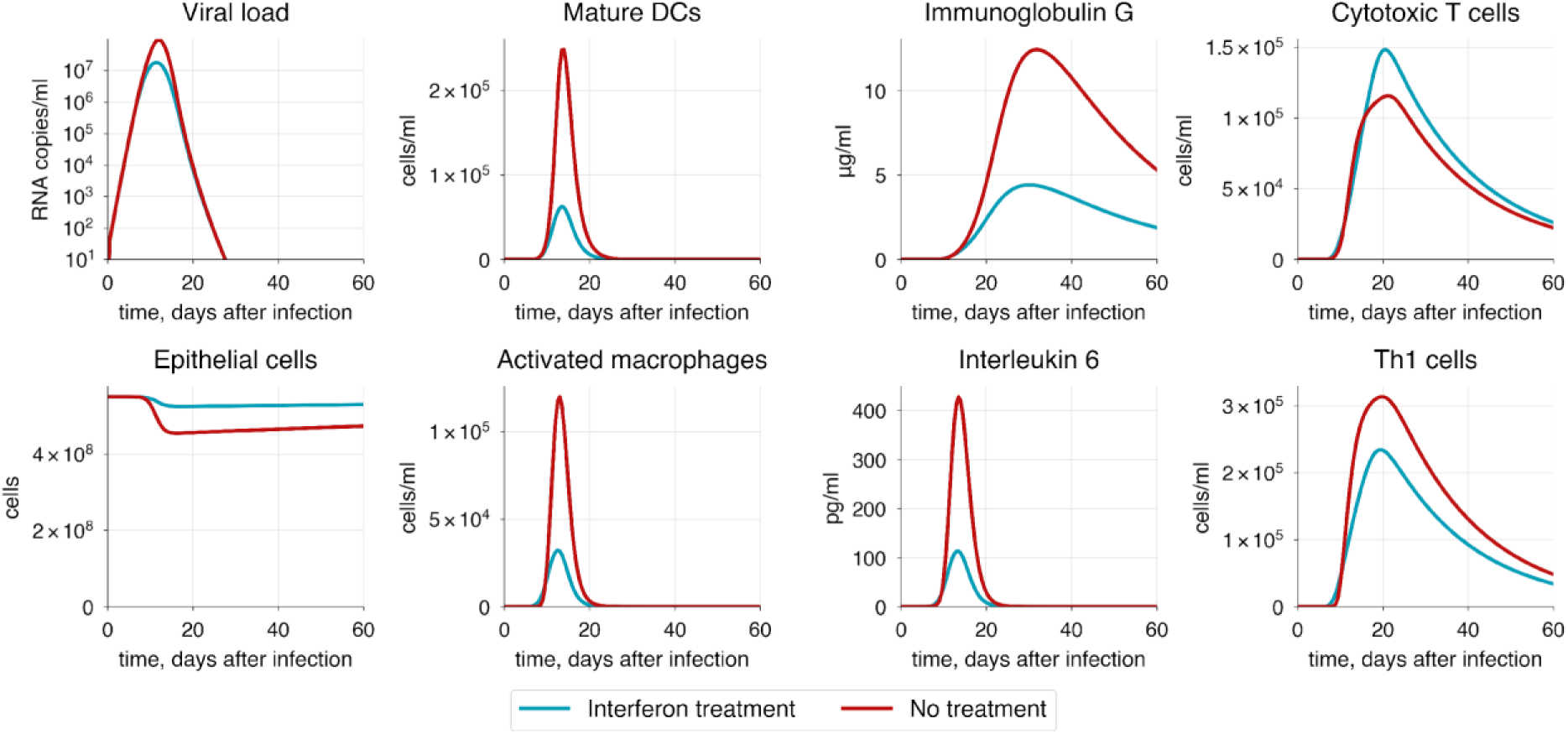
The baseline model solution with interferon administration. Interferon is administered at a concentration of 2000 pg/ml daily for five days post-symptom onset.

Another critical target for antiviral drugs is the suppression of the viral replication process. Examples of such drugs include amodiaquine, atovaquone, bedaquiline and others [177]. Using the developed model, we studied the effect of daily drug injections on inhibiting viral replication within infected cells, simulating the drug administration. We assumed that the replication parameter 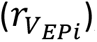 changes dynamically during the course of treatment before returning to its baseline value, reflecting the temporary nature of the drug’s action. During drug administration we observed a significant reduction in viral load and damage to epithelial cells, demonstrating the high efficacy of these drugs and their potential for treating COVID-19 (Figure 16).

**Figure 16.**
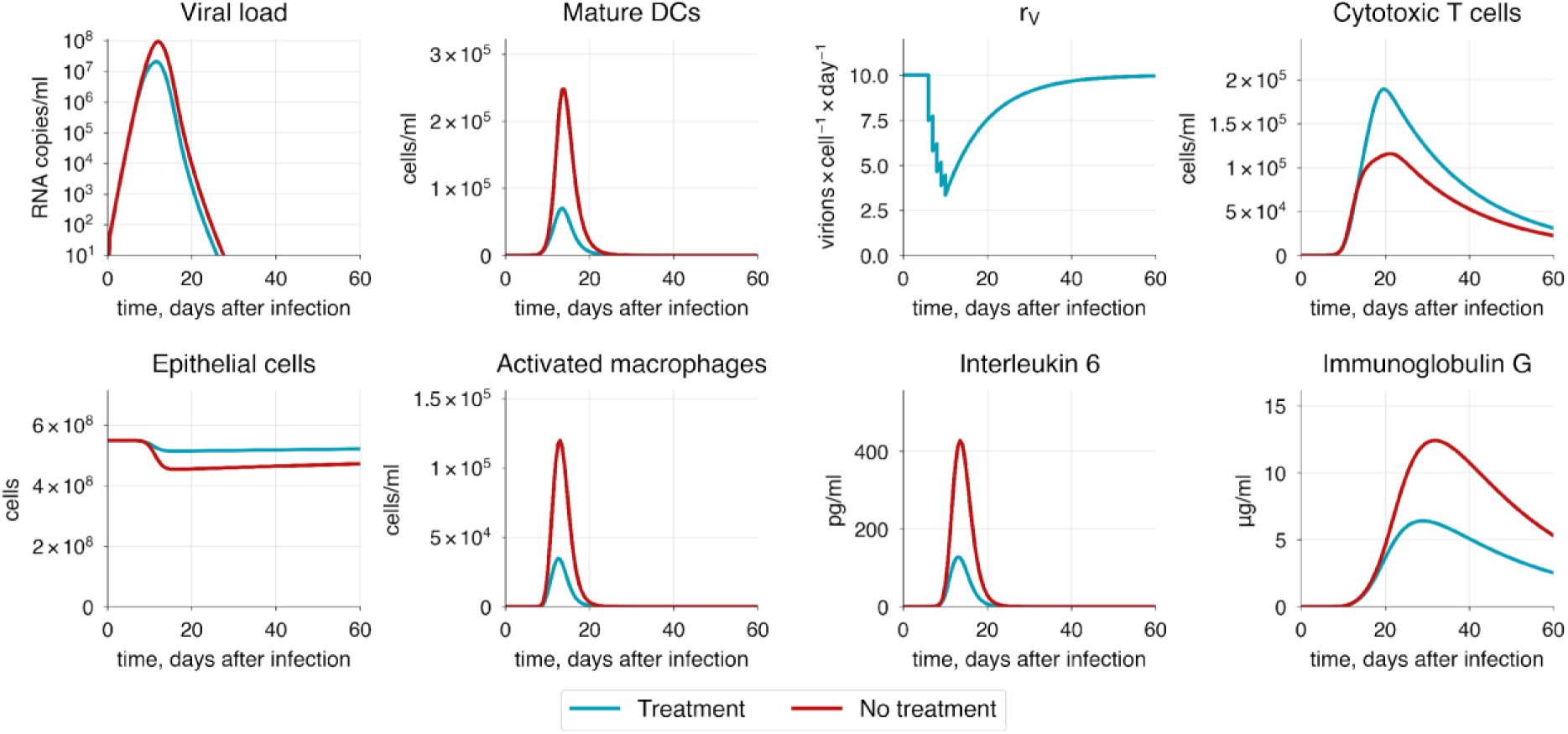
The baseline model solution with inhibited viral replication. Drug is administered daily for five days post-symptom onset.

## Conclusions

Herein, we present a developed modular model of the human immune response to SARS-CoV-2 infection. The model structure consists of two main parts: the upper airways (including the nasal cavity, oral cavity, and pharynx) and the lower airways (including the trachea, bronchial tree, and lungs). Each structural module is connected to a draining lymph node compartment, in overall forming four interconnected modules. The model incorporates innate immunity, involving macrophages and dendritic cells, and adaptive immunity, comprising humoral (antibodies) and cellular (CTLs, Th1, and Tfh cells) components. Cellular interactions in the model are facilitated by direct contacts, such as antigen presentation, or by cytokine signaling via IL-2, IL-6, IL-12, and IFNγ.

To estimate the model parameters, we used time series clinical data on moderate disease progression, including concentrations of IL-6, CD4+ and CD8+ T cells, IgA, IgM, IgG, and viral load in both the upper and lower respiratory tracts. Additionally, we considered age-related changes in the immune system to model severe and critical disease states. Severity state was determined by the extent of lung tissue damage, elevated levels of IL-6 and viral load, as well as by the observed reasonable decline in concentrations of neutralizing antibodies and CTLs. To validate the model, we conducted a series of *in silico* experiments focusing on key processes: infectivity and evasion mechanisms of SARS-CoV-2, cytotoxicity of T cells and antibody neutralization activity, as well as innate immune response delay and hyperactivation. Finally, we simulated HIV/SARS-CoV-2 coinfection and demonstrated the efficacy of several therapeutic strategies for treatment of COVID-19 patients including interferon administration.

Aging is recognized as the most fundamental risk factor for developing severe COVID-19, potentially leading to a fatal outcome. To investigate its impact, we defined a set of parameters (ε) representing key age-related processes, including the number of naïve lymphocytes, their proliferation and differentiation rates, and the efficacy of innate immunity activation. Simulation results show that a more than 30% overall decrease in the immune response efficacy leads to critical COVID-19 progression, characterized by significant alveolar epithelium damage, higher viral load, and an approximately 40% increase in viral shedding duration compared to moderate progression. Additionally, we incorporated the cytokine storm mechanism, a hallmark of critical COVID-19 cases often requiring mechanical ventilation. It is characterized by the excessive release of pro-inflammatory cytokines by innate immune cells such as macrophages, with Interleukin 6 being a key contributor. While IL-6 promotes B cell development, its elevated level inhibits proper T cell formation and suppresses the cytotoxic activity of CTLs. These findings highlight the complex interplay between aging and immune response in the progression of severe COVID-19.

To investigate the contribution of the innate immune response to the infection, we separately examined the hyperactivation of macrophages and the delay in dendritic cell maturation and migration. These factors contribute to a worse disease outcome, significantly increasing the likelihood of a cytokine storm, as confirmed by substantially elevated IL-6 levels (>1000 pg/ml). Furthermore, a sensitivity analysis of the built model highlights the important role of dendritic cell migration, showing high sensitivity to delay in IDC migration to lymph nodes. This underscores the significance of innate immunity, especially in activating the adaptive immune response.

Patients with chronic diseases like HIV are at higher risk of developing respiratory illnesses such as COVID-19 or influenza, which can lead to co-infection with multiple pathogens. Although modeling coinfection is challenging due to its complexity, we can simulate the co-infection by focusing on the main consequences of chronic infection. For instance, in the case of HIV, the key change is the depletion of CD4+ T cells, which severely impairs the immune system’s ability to mount a strong response. The reconstructed model predicts significantly worse disease progression in these patients. Specifically, a reduction in CD4+ T cells by more than 50% leads to a critical state of COVID-19, causing serious alveolar epithelium damage and thus requiring hospitalization and mechanical ventilation.

Currently, many approaches to COVID-19 therapy have been developed, with vaccination being the most common. Although vaccination primarily serves as a preventive measure, there is a need for effective treatment strategies to mitigate the severity of infection in patients who are already experiencing symptoms. One promising approach is interferon administration, which may improve the condition of critically ill patients and prevent the development of more severe progression of COVID-19. To verify the effect of interferon administration on the immune response, we focused on protocols used by healthcare providers that involve daily injections of interferon gamma for 5-6 days, starting immediately after symptom onset. According to the model predictions, the treatment not only reduces viral load and alveolar epithelium damage by more than 50%, but also raises baseline interferon levels even after the treatment ends, leading to a milder course of COVID-19 and ensuring faster recovery.

The proposed modular model primarily incorporates viral load dynamics with host immune response including both arms of the immune system. We aim to further extend the model by incorporating cytokines and T helper cells into the upper airways compartment and via an explicit addition of the blood system module for a more accurate representation of transport processes within the body. The adaptive immune response is inherently complex and regulated by numerous feedback mechanisms, with T regulatory cells playing a crucial role. Consideration of these cells is also a key aspect of the model development. The ultimate goal of the modular approach used in the study is to create a comprehensive multi-compartmental model that accurately reflects both innate and adaptive immune responses to various pathogens. Consequently, we believe that the constructed model framework can also be applied to study other pathogens like the influenza virus or an entirely novel agent, and to predict clinical outcomes in COVID-19 patients depending on their age, immune status and given different therapeutic treatments.

## Author Contributions

Conceptualization, M.I.M. and I.R.A.; methodology, M.I.M., I.R.A. and F.A.K.; software, F.A.K.; validation, M.I.M. and I.R.A.; data curation, M.I.M.; writing—review and editing, M.I.M. and I.R.A.; funding acquisition, F.A.K. All authors have read and agreed to the published version of the manuscript.

## Data Availability Statement

The multi-scale mathematical model of the immune response to SARS-CoV-2 infection as well as all simulation results described in the manuscript are available through the web interface of BioUML software at GitLab project: https://gitlab.sirius-web.org/diploms/modular-immune-system. It is worth noting that simulation results are presented via Jupyter Notebooks for the independent validation of the model simulations and further numerical analysis. Furthermore, to simplify the model simulations we have added the widget to the Notebook for different simulation scenarios (see readme in the GitLab project).

